# An integrative structural model of the full-length gp16 ATPase in bacteriophage phi29 DNA packaging motor

**DOI:** 10.1101/2020.07.20.213124

**Authors:** Abdullah F.U.H. Saeed, Chun Chan, Hongxin Guan, Bing Gong, Peixuan Guo, Xiaolin Cheng, Songying Ouyang

**Affiliations:** The Key Laboratory of Innate Immune Biology of Fujian Province, Provincial University Key Laboratory of Cellular Stress Response and Metabolic Regulation, Biomedical Research Center of South China, Key Laboratory of OptoElectronic Science and Technology for Medicine of Ministry of Education, College of Life Sciences, Fujian Normal University, Fuzhou, 350117, China; Laboratory for Marine Biology and Biotechnology, Pilot National Laboratory for Marine Science and Technology (Qingdao), Qingdao 266237, China; College of Chemistry and Materials Science, Fujian Normal University, Fuzhou 350117, China; College of Pharmacy, The Ohio State University, Columbus, OH, USA; Guangxi Key Laboratory of Marine Disaster in the Beibu Gulf, Beibu Gulf University, Qinzhou, 535000, China; Center for RNA Nanobiotechnology and Nanomedicine; College of Medicine; Dorothy M. Davis Heart and Lung Research Institute; Comprehensive Cancer Center, The Ohio State University, Columbus, OH, USA; Biophysics Graduate Program; Translational Data Analytics Institute, The Ohio State University, Columbus, OH, USA

**Keywords:** DNA packaging motor, ATPase, crystallography, integrative modeling

## Abstract

Biological motors, ubiquitous in living systems, convert chemical energy into different kinds of mechanical motions critical to cellular functions. Most of these biomotors belong to a group of enzymes known as ATPases, which adopt a multi-subunit ring-shaped structure and hydrolyze adenosine triphosphate (ATP) to generate forces. The gene product 16 (gp16), an ATPase in bacteriophage □29, is among the most powerful biomotors known. It can overcome substantial resisting forces from entropic, electrostatic, and DNA bending sources to package double-stranded DNA (dsDNA) into a preformed protein shell (procapsid). Despite numerous studies of the □29 packaging mechanism, a structure of the full-length gp16 is still lacking, let alone that of the packaging motor complex that includes two additional molecular components: a connector gp10 protein and a prohead RNA (pRNA). Here we report the crystal structure of the C-terminal domain of gp16 (gp16-CTD). Structure-based alignment of gp16-CTD with related RNase H-like nuclease domains revealed a nucleic acid binding surface in gp16-CTD, whereas no nuclease activity has been detected for gp16. Subsequent molecular dynamics (MD) simulations showed that this nucleic acid binding surface is likely essential for pRNA binding. Furthermore, our simulations of a full-length gp16 structural model highlighted a dynamic interplay between the N-terminal domain (NTD) and CTD of gp16, which may play a role in driving DNA movement into the procapsid, providing structural support to the previously proposed inchworm model. Lastly, we assembled an atomic structural model of the complete □29 dsDNA packaging motor complex by integrating structural and experimental data from multiple sources. Collectively, our findings provided a refined inchworm-revolution model for dsDNA translocation in bacteriophage □29 and suggested how the individual domains of gp16 work together to power such translocation.

**ABSTRACT (SHORT):** Biological motors, ubiquitous in living systems, convert chemical energy into different kinds of mechanical motions critical to cellular functions. The gene product 16 (gp16) in bacteriophage □29 is among the most powerful biomotors known, which adopts a multi-subunit ring-shaped structure and hydrolyzes ATP to package double-stranded DNA (dsDNA) into a preformed procapsid. Here we report the crystal structure of the C-terminal domain of gp16 (gp16-CTD). Structure-based alignment and molecular dynamics (MD) simulations revealed an essential binding surface of gp16-CTD for prohead RNA (pRNA), a unique component of the motor complex. Furthermore, our simulations highlighted a dynamic interplay between the N-terminal domain (NTD) and CTD of gp16, which may play a role in driving DNA movement into the procapsid. Lastly, we assembled an atomic structural model of the complete □29 dsDNA packaging motor complex by integrating structural and experimental data from multiple sources. Collectively, our findings provided a refined inchworm-revolution model for dsDNA translocation in bacteriophage □29 and suggested how the individual domains of gp16 work together to power such translocation.

## Introduction

During the late stage in morphogenesis, double-stranded DNA (dsDNA) viruses package their genomes into a preformed protein shell (procapsid). Genome encapsidation is an extremely unfavorable process, both entropically and enthalpically, which is accomplished by a viral DNA packaging motor. In 1986, this DNA packaging motor (also known as gene product 16, gp16) was first reported to be an ATPase ^1^ that hydrolyzes ATP to drive DNA packaging. Titration assays using a thin-layer chromatography for radiative ATP quantification concluded that one molecule of ATP was consumed to package every 2 DNA base pairs ^1^. A subsequent study showed that ATP hydrolysis by gp16 was both prohead and DNA-gp3 dependent^2^. The same study also identified the ATP-binding (Walker residues^3^) and potential magnesium-binding domains in gp16, which were also present in other phage packaging motors, such as gpA of λ, gp19 of T7 and gp17 of T4.

ATPase gp16, residing at a unique vertex of the viral shell, belongs to a class of ubiquitous ring-shaped nucleoside triphosphate (NTP)-dependent molecular motors. These motors are powerful enough to package DNA against extremely high internal pressure (tens of atmospheres) established by densely packed negatively charged DNA inside the shell. As such, there have been extensive efforts in engineering viral packaging motors as a functional nanodevice for the delivery of nucleic acid therapeutics. Among the various motors, bacteriophage □29 has been developed into a highly efficient *in vitro* system ^1^ and therefore served as an excellent model system for structural and mechanistic studies of biomotors ^4^ as well as nanotechnological development ^5,6^.

Viral packaging motors can be classified into two distinct families: the terminase family and the HerA/FtsK-type ATPases. Although all known dsDNA packaging ATPases belong to the P-loop NTPase fold under the ASCE (additional strand conserved E (glutamate)) division, sequence and structural analyses were able to distinguish bacteriophage □29 from the terminase-family viruses and place it as a divergent version of the HerA/FtsK superfamily ^7^. A number of viral packaging motor structures have been resolved, namely the large terminase (TerL) proteins, of the terminase family, such as bacteriophage T4 gp17 ^8^, Sf6 gp2 ^9^, D6E TerL protein ^10^, thermophilic phage P74-26 TerL ^11,12^, and herpesvirus herpes simplex virus 1 (HSV1) pUL15 ^13^. The TerL proteins are the catalytic engine of DNA packaging motors, which harbor two enzymatic functionalities: an adenosine triphosphatase (ATPase) that consumes ATP to translocate DNA, and an endonuclease that cleaves genome concatemers at both initiation and termination steps of packaging. By contrast, □29 lacks the cleavage (nuclease) functionality and packages its unit-length genome covalently attached by a recognition protein ^1,14^. The gp16 structure has proven to be extremely challenging to resolve. Only until very recently, Mao *et al*. reported an X-ray crystallographic structure of the N-terminal domain (ATPase domain) of gp16 and a pseudo-atomic structure of the functional motor complex that also included the connector protein and the prohead RNA (pRNA) besides gp16 by docking the various component structures into low-resolution (12 □) cryo-EM maps ^4^.

Partially due to a limited atomistic view of the viral packaging motor proteins, the oligomeric state of the individual motor components has been debated for decades. The debate initiated in the stoichiometry of pRNA, which is a unique component of the □29 packaging motor complex ^15^. While early biochemical assays supported a hexameric ring structure of pRNA ^16,17^, Cao *et al*. later confirmed this result but also showed an alternative pentameric pRNA ring structure from asymmetric reconstruction of the cryo-EM data ^18,19^. More recent cryo-EM data have also favored a pentameric structure ^4,20^. The same mystery carried over to the gp16 ATPase where low-resolution cryo-EM reconstructions slightly favored 5-fold symmetry averaged structures ^21–23^, whereas a large body of biochemical data supported a hexameric ring structure ^16,24,25^. Similarly in bacteriophage T4 and T7, atomic structures of the packaging motor proteins seemed to fit both 5- and 6-fold symmetry averaged low-resolution (16-32 □) cryo-EM densities ^8,26^. All these controversies are rooted in the lack of a high-resolution structure of the □29 packaging motor complex. Recently, Yang *et al*. have determined the first atomic structures of a herpesvirus terminase complex in both apo and ATP mimic-bound states and provided a reconciling structural view of this class of viral dsDNA packaging motors ^13^.

The □29 gp16 ATPase is a 39-kDa 332-amino-acid protein consisting of two domains connected by a flexible linker. The N-terminal domain (gp16-NTD, residues 1-208) is approximately 200 amino acid long and contains the conserved ASCE ATPase domain. The C-terminal domain (gp16-CTD, residues 228-332) contains about 100 residues, but little is known about its function. The X-ray crystallographic structure of gp16-NTD (residues 4-197) has been determined previously ^4^. However, crystallization of the □29 full-length gp16 (FL-gp16) has been difficult, so an atomic structure of FL-gp16 is still lacking, leaving a knowledge gap of how the two domains are spatially organized and interact with each other as well as the pRNA and the connector. Here, we reported an X-ray crystallographic structure of gp16-CTD at 2.3 Å resolution. By aligning this structure to those of its counterparts in the TerL complexes, we identified possible pRNA or DNA binding surfaces of gp16-CTD. The availability of an atomic structure of gp16-CTD together with existing structural information of the other motor components has also enabled us to build the first atomic structural model of FL-gp16. Our molecular dynamics (MD) simulations of FL-gp16 further highlighted that the two domains are free to translate and reorient relative to each other, resulting in a wide range of conformations. We finally employed an integrative structural modeling strategy and MD simulations to assemble a pseudo-atomic model of the □29 packaging motor complex, shedding light on a critical coordinative role of gp16-CTD in motor assembly and operation, and the mechanisms by which the ring-shaped ATPases translocate their substrates.

## Results and discussion

### Structure of the □29 gp16 C-terminal domain

The gp16-CTD structure displays a compressed football shape with dimensions of 18 Å × 20 Å × 12 Å. A central five-stranded β sheet (β1-β5), with mixed parallel and antiparallel strands, is flanked on one side by two α helices through hydrophobic interactions (Figure 1A). An extended Loop 1 (L1) that connects β3 and β4 covers the other side of the β sheet by crossing over the β1 and β2 strands. The strand order in the central sheet is 4, 1, 2, 3, and 5. On one lateral edge of the central β-sheet, Loop 2 (L2), formed between β4 and α1, holds a twisted β-hairpin. On the other edge, β3, β5, and the N-terminal part of α2 surround a cleft that typically harbors the conserved active site residues (dotted circle in Figure 1A) in the RNase H/resolvase/integrase superfamily enzymes.

**Figure 1.**
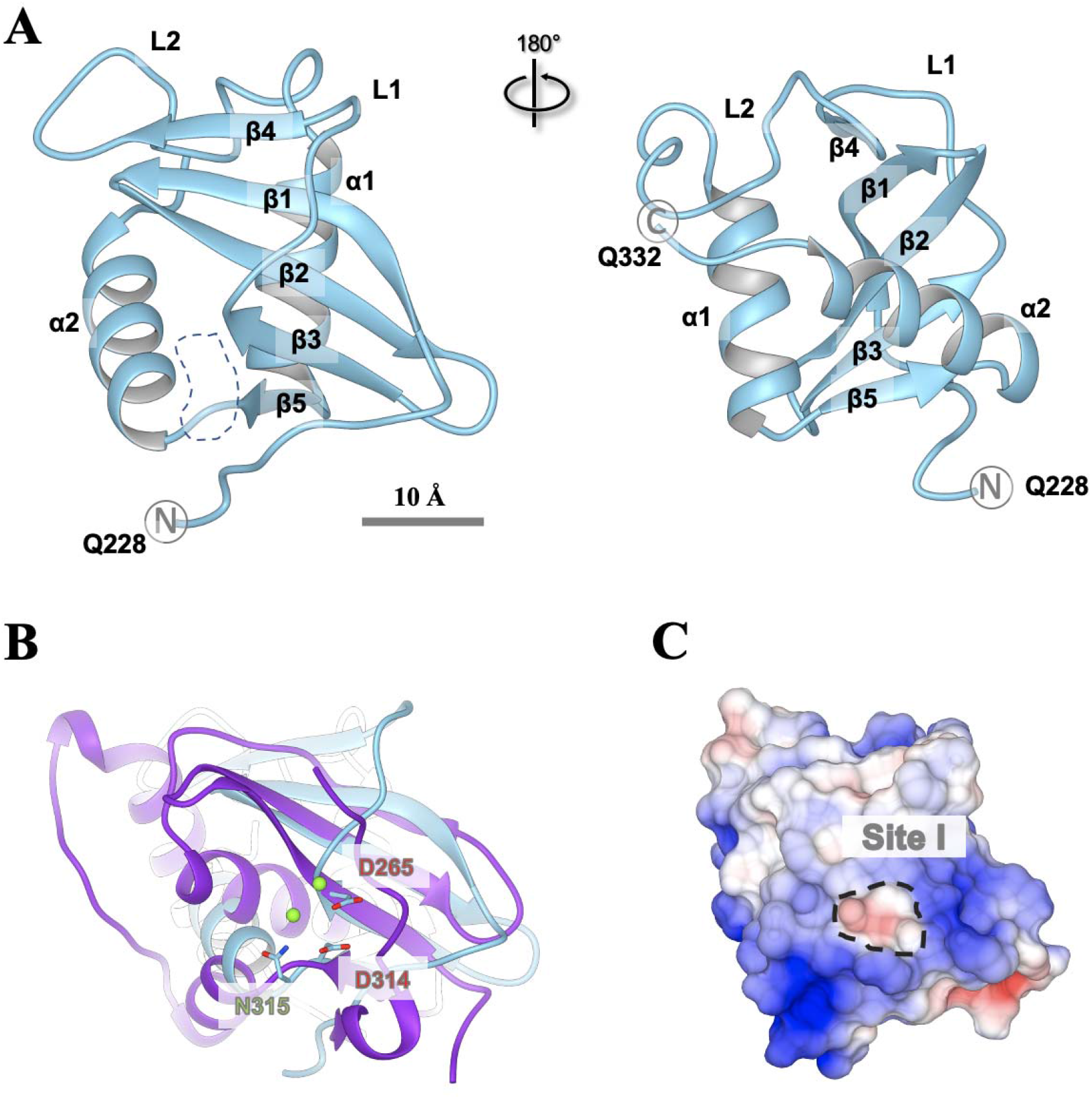
X-ray crystallographic structure of the gp16 C-terminal domain (gp16-CTD). (A) Cartoon diagrams of gp16-CTD. α helices and β strands are labeled and numbered. The analogue active site (Site I) is circled by dotted line. (B) Superposition of gp16-CTD (colored in cyan) with RNase H1 (colored in purple; PDB: 1ZBI). Charged residues at Site I of gp16-CTD are labeled and shown as sticks. Magnesium ions from the crystallographic structure of RNase H1 are shown in green spheres. (C) Electrostatic surface of gp16-CTD. Positive and negative charges are colored in blue and red, respectively. Site I is circled by dotted line.

The gp16-CTD structure resembles the RNase H fold, with the greatest similarity to RNase H1 (Figure 1B). This topological feature is conserved among all CTDs of the packaging ATPases in bacteriophages and herpesviruses. Despite low sequence identity (< 15%), the central β-sheet of gp16-CTD superimposes well its counterparts in other terminase large subunits (TerL), such as T4 gp17, SPP1 gp2 and HSV1 pUL15 (Figure S1). Interestingly, although no nuclease activity of gp16-CTD has been reported to date, conserved RNases H acidic residues D265 and D314 are found on one edge of the central β-sheet where two parallel β-strands (β3 and β5) orient in a fork-like manner (Figure 1B). D265 is located at the end of β3 while D314 at the end of β5, forming a small electronegative site (Site I) on the protein surface (Figure 1C). Conventionally, three conserved acidic residues constitute the active center in the RNase H family of nucleases; for example, they are three aspartic acid residues in RNase H ^27,28^. A two-metal-ion mechanism was proposed for the catalytic function of this family of nucleases^29^. Two metal ions (Mg^2+^ or Mn^2+^) were found to be coordinated by the three conserved acidic residues in the structure of RNase H complexed with an RNA/DNA hybrid ^27^. These acidic residues are structurally conserved among the RNase H-like nuclease domains in other phages as well, such as T4 gp17 ^8^. However, superimposition of gp16-CTD onto the RNase H shows that the third catalytic aspartic acid is replaced by an asparagine in gp16-CTD (Figure 1B). Although the two aspartic acids are conserved, the lack of the third Asp may account for its deficiency in nuclease activity. The retention of this mysterious catalytic site is a clear example of divergent evolution, but the new role of this heritage site and the evolution of its role adapted to its motor functional requirement remain elusive.

### Nucleic acid binding surfaces on gp16-CTD

Despite the conservation of the overall topology and the active site residues, there is very little sequence conservation in the RNase H-like domains between bacteriophages and herpesviruses. This lack of conservation suggests that different viruses package their genomes differently. We hypothesized that although the ϕ29 DNA packaging ATPase is shown not to possess the nuclease activity, the RNase H-like structure may still be preserved and adapted for nucleic acid binding. Nevertheless, due to the low sequence homology, determining how gp16-CTD interacts with various types of nucleic acids, such as pRNA or dsDNA substrate, has been particularly challenging.

Similar to its counterparts in other RNase H-like nuclease domains, Site I of gp16-CTD constitutes a shallow groove, with its edge lined by positively charged residues, K233, R234, K236, K239, H268, R312, and R319. The groove bottom is formed by β2 and β3, which contribute a tryptophan W254 and a tyrosine Y263, respectively, to make stacking interactions with nucleic acid bases (Figure 2A). The N-terminus of gp16-CTD, which contains three nearly consecutive basic residues K233, R234, and K236, divides the site into a major and a minor binding grooves (colored in yellow and magenta, respectively in Figure 2B). To explore the conformational dynamics of Site I, we performed MD simulations of gp16-CTD in aqueous solution. The root-mean-square-fluctuation (RMSF) profile indicates that the N-terminus (residues 228-238) of gp16-CTD is relatively rigid, in agreement with the low B-factor values for this region in the crystallographic structure (Figure S2). The rigid N-terminus excludes the possibility of forming a larger binding surface by large displacements of the N-terminus via thermal fluctuation. The solvent-accessible surface areas (SASAs) of both the major and minor binding grooves are invariant throughout the MD simulations (Figure S3), implying the importance of the basic patch K233, R234, and K236 for nucleic acid binding.

**Figure 2.**
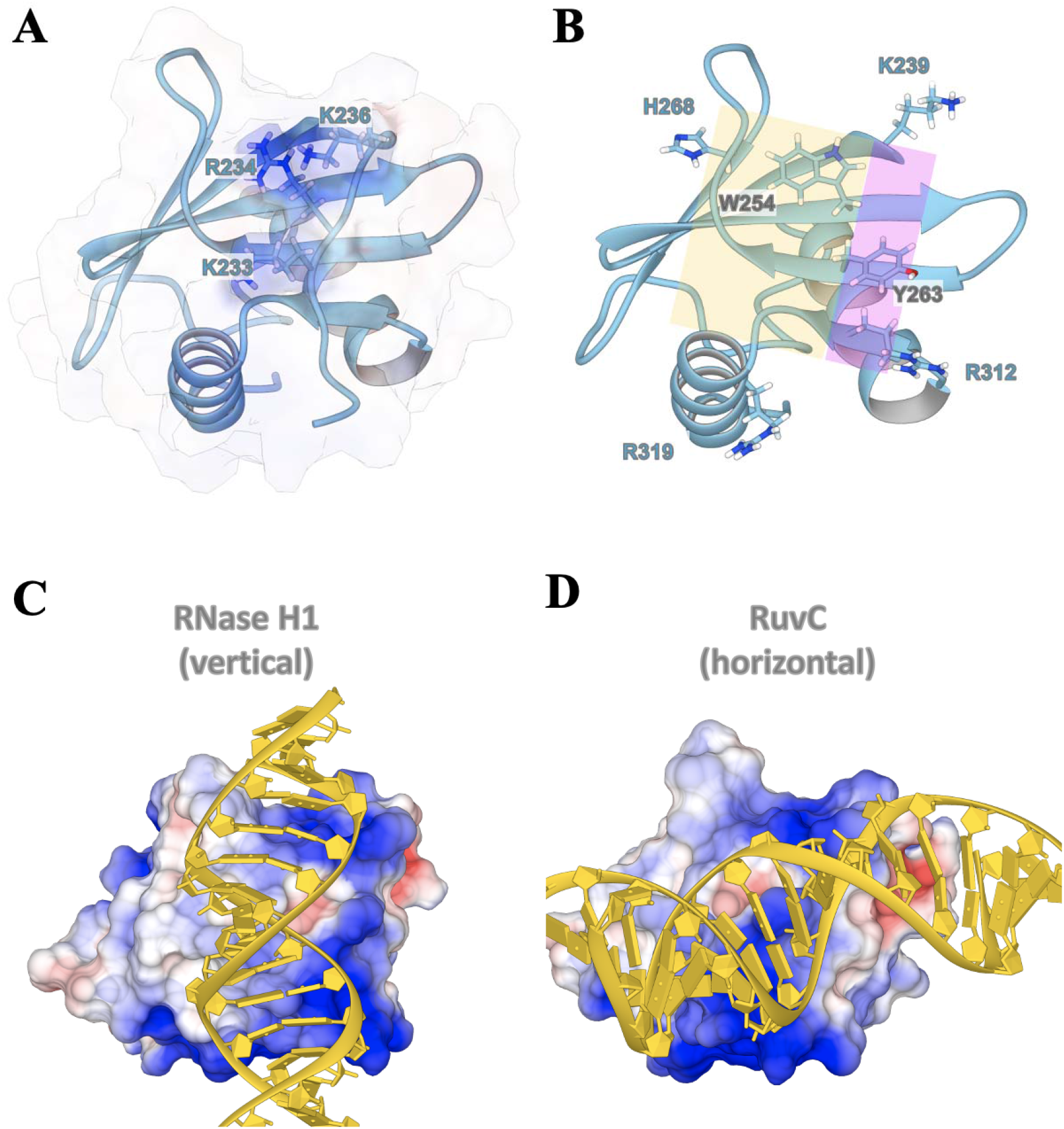
The analogous active site suggests a nucleic acid binding surface for gp16-CTD. (A) Charged residues on the N-terminus of gp16-CTD form a positively charged surface. Charged residues are labeled and shown as sticks. (B) Two binding surfaces separated by the N-terminus of gp16-CTD. The major and the minor binding surfaces are colored in yellow and magenta, respectively. Charged residues surrounding the surface and hydrophobic residues at the center of the surface are labeled and shown as sticks. For clarity, the N-terminus is not shown. (C) Superposition of gp16-CTD with two RNase H-like domains suggested two potential DNA binding orientations (vertical and horizontal) near Site I of gp16-CTD.

To uncover how Site I may accommodate nucleic acids, we compared the gp16-CTD structure to those of several distantly related nucleases bound with substrates, which suggested two possible DNA binding orientations in the groove. First, superposition of the RNase H/RNA-DNA structure with gp16-CTD placed the DNA helix on the gp16-CTD surface, extending from the N-terminus of the gp16-CTD to the C-terminal end of L1 (Figure 2C). We designate this orientation as the vertical orientation. In contrast, superposition with a very recent structure of the *Thermus thermopilus* RuvC/Holiday junction complex placed the DNA helix in an orthogonal orientation, denoted as the horizontal orientation (Figure 2D). Similar DNA orientations were identified for TerL of thermophilic phage P74-26 also by superposition with RNase H and RuvC ^12^. However, we note a crucial difference between the two comparisons. In P74-26, a β-hairpin in the viral TerL proteins clashed with the vertically placed DNA helix, whereas this β-hairpin is absent in gp16-CTD and all other known members of the RNase H family ^28,30,31^. On the other hand, the L1 loop (residue 266-274) in gp16-CTD is significantly shorter than the β-hairpin in the TerL proteins, possibly because a flexible β-hairpin that plays an auto-regulatory role in nuclease function is not required by gp16-CTD. Our MD simulations confirmed that L1 has negligible flexibility and would minimally interfere with the DNA binding surface of gp16-CTD (Figure S2). Thus, no clashes were found for either vertical or horizontal binding of DNA on gp16-CTD. Furthermore, a positively charged N-terminus of gp16-CTD can insert into the DNA major groove, stabilizing DNA binding in both vertical and horizontal orientations. These results suggest that gp16-CTD may have multiple nucleic acid binding modes. Importantly, our identified nucleic acid binding surfaces on gp16-CTD are not limited to DNA. Instead, gp16-CTD may also interact with pRNA through these surfaces, which could shed light on the essential roles of pRNA in motor assembly and operation ^15^.

Besides Site I, gp16-CTD contains another surface site enriched in positively charged residues, hereafter denoted as Site II. This site is mainly formed by two basic patches near the N-terminus of helix α1 and the C-terminus of helix α2 (Figure S4). Our MD simulations showed that despite the favorable electrostatic interactions, L2 sandwiched between the two basic patches is highly flexible, consistent with the large B-factors observed in the crystallographic structure (Figure S2). The highly dynamic L2 can kinetically weaken the interaction of nucleic acids with gp16-CTD, making Site II a likely transient site for substrate binding.

### The linker domain connecting NTD and CTD of gp16

The NTD and CTD domains of gp16 are connected by a linker domain that may play an important role in mediating the domain interactions and thus shaping the structure and function of the entire gp16 motor. Additionally, the linker appears to make direct contact with helix α2 that constitutes the nucleic acid binding Site I in gp16-CTD. Thus, the linker may also be involved in interacting with pRNA or DNA substrate during packaging. The structure of the gp16 linker domain (residues 198-227) was not resolved in either previous gp16-NTD structure ^4^ or our current gp16-CTD structure. The gp16 linker is significantly shorter, with only 30 residues as compared to 90-150 residues in other viral ATPases. In T4 gp17, the linker was identified as a subdomain of NTD and made extensive interactions with the substrate DNA ^8^. In phage Sf6 gp2, the linker together with the nuclease domain formed the DNA binding surface ^9^.

We performed MD simulations on the gp16 linker domain starting from an extended chain conformation. The results showed that the linker domain remained as a flexible chain lacking a secondary structure throughout the simulation, in agreement with the secondary structure prediction using PsiPred ^32^ (Figure S5). Given that electron densities of both NTD and CTD were resolved in the cryo-EM data ^4^, the linker domain is believed to occupy the space between the two domains, but its electron density map is blurred due to its high flexibility. This inherent disordered nature of the linker also explains why it was difficult to resolve the structure in crystallography or cryo-EM.

### Dynamic interactions between NTD and CTD of gp16

To begin understanding the roles of the individual gp16 domains in genome packaging, we have used an integrative structural modeling approach to build an atomic structure of the full-length gp16 (FL-gp16). A key challenge in building an FL-gp16 structural model is to determine the relative position and orientation of the two terminal domains. To do this, we first compared gp16-NTD to the ATPase domains of other viral motor proteins. Gp16-NTD displays a canonical ASCE fold commonly found in ATPases such as F1-ATPase^33^, DNA/RNA helicases or translocases^34–36^, and viral packaging motors^4,7,26,37^. Like other members of the family, the gp16-NTD core contains a five-stranded, parallel β-sheet sandwiched between four α-helices. The structure of gp16-NTD closely resembles that of the TerL ATPase domain ^4^. However, based on the location of the putative arginine finger (R146) and the lack of nuclease function in gp16-CTD, gp16 was also proposed to belong to the HerA/FtsK branch of the ASCE superfamily ^7^. Despite the debate, we chose to compare gp16 with the most recently resolved cryo-EM structure of Herpes Simplex Virus (HSV1) ATPase/terminase pUL15 ^13^. pUL15 is the only TerL protein resolved in its functional oligomeric state, whereas crystal structures of several full-length viral large terminase subunits, such as T4 gp17 and Sf6 gp2, have been available for years ^8,9^.

We determined the relative orientation of gp16-NTD in a functional motor complex by comparing it to that of pUL15 with respect to its DNA substrate. To assess the feasibility of this approach, we determined the structural homology between the ATPase domains of gp16 and pUL15 by comparing gp16 to several other viral packaging ATPases (Table S1). Upon structural superposition, the ATPase domain of gp16 aligned considerably well with that of pUL15 despite their distant relation in the viral family. A homology modelled structure of gp16-NTD based on pUL15 showed a low root-mean-square-deviation (RMSD) for a large number of Cα atoms when aligned with its crystallographic structure (PDB: 5HD9; Table S1).

We then explored the relative position and orientation of CTDs in several full-length monomer motor proteins by superimposing their structures using the ATPase domains as a reference (Figure 3). Interestingly, we found significant variations in both NTD-CTD separation and orientation. In pUL15, the linker occupies the space between NTD and CTD that make no contact. The catalytic site of the nuclease domain in pUL15 opens towards the adjacent subunit instead of the substrate DNA, suggesting substrate-based auto-inhibition of the nuclease activity during translocation ^13^. By contrast, the crystallographic structures of the motor proteins T4 gp17 ^8^ and Sf6 gp2 ^12^ both show considerable NTD-CTD interactions. The catalytic residues on CTD are either covered by the linker domain in T4 or the adjacent subunit in Sf6, also suggesting the inaccessibility of the nuclease site to DNA during translocation. Evidently, conformational changes of CTD are required for the motor proteins to switch between a translocation and cleavage mode, but the molecular details of these conformational changes remain largely unknown. A variety of NTD-CTD separations and CTD orientations observed in recent crystal and cryo-EM structures suggest the high plasticity of the viral motor proteins, which can be partially attributed to the intrinsically disordered (unstructured) linker domain.

**Figure 3.**
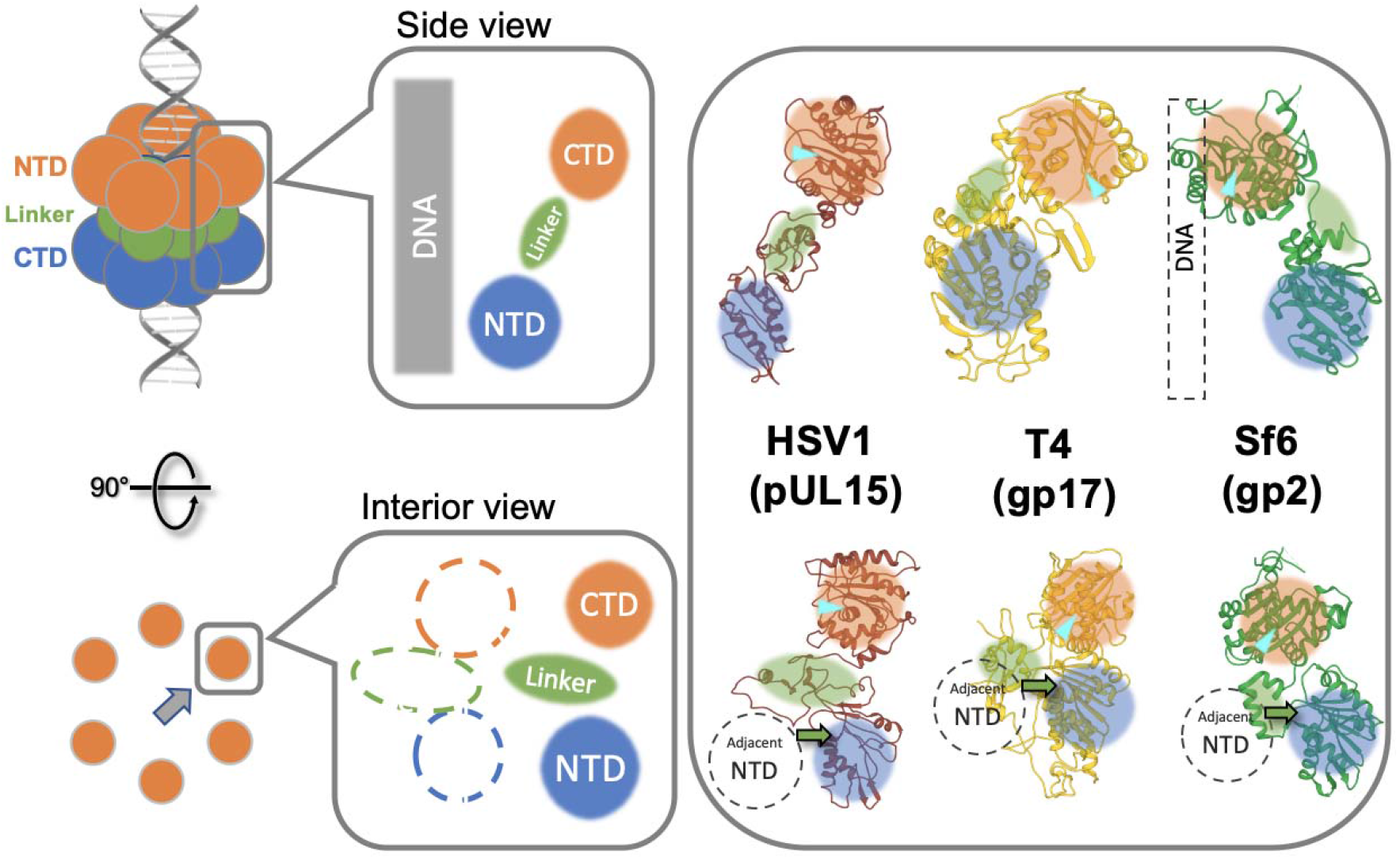
Structural superposition of several viral packaging motor proteins according to their ATPase domains (NTD). CTD, linker and NTD are indicated by orange, green and blue circles, respectively. The nuclease active sites on CTD are indicated by cyan triangles. Two views are shown. The side view (above) refers to a cross-section view along the DNA translocation axis. The interior view (below) refers to viewing from the interior of the central channel along the radial axis.

To probe the dynamic interactions between gp16-CTD and gp16-NTD, we performed MD simulations on an FL-gp16 model (Figure 4A; Movie S1) that was built by combining the crystallographic structures of gp16-CTD and gp16-NTD, and an MD-equilibrated gp16 linker (details in Methodology). To avoid biasing towards the initial conformations due to insufficient sampling, we have performed eight independent simulations started from eight different FL-gp16 models with CTD in four different orientations and large separation between CTD and NTD (center-of-mass (COM) distance ~60 Å).

**Figure 4.**
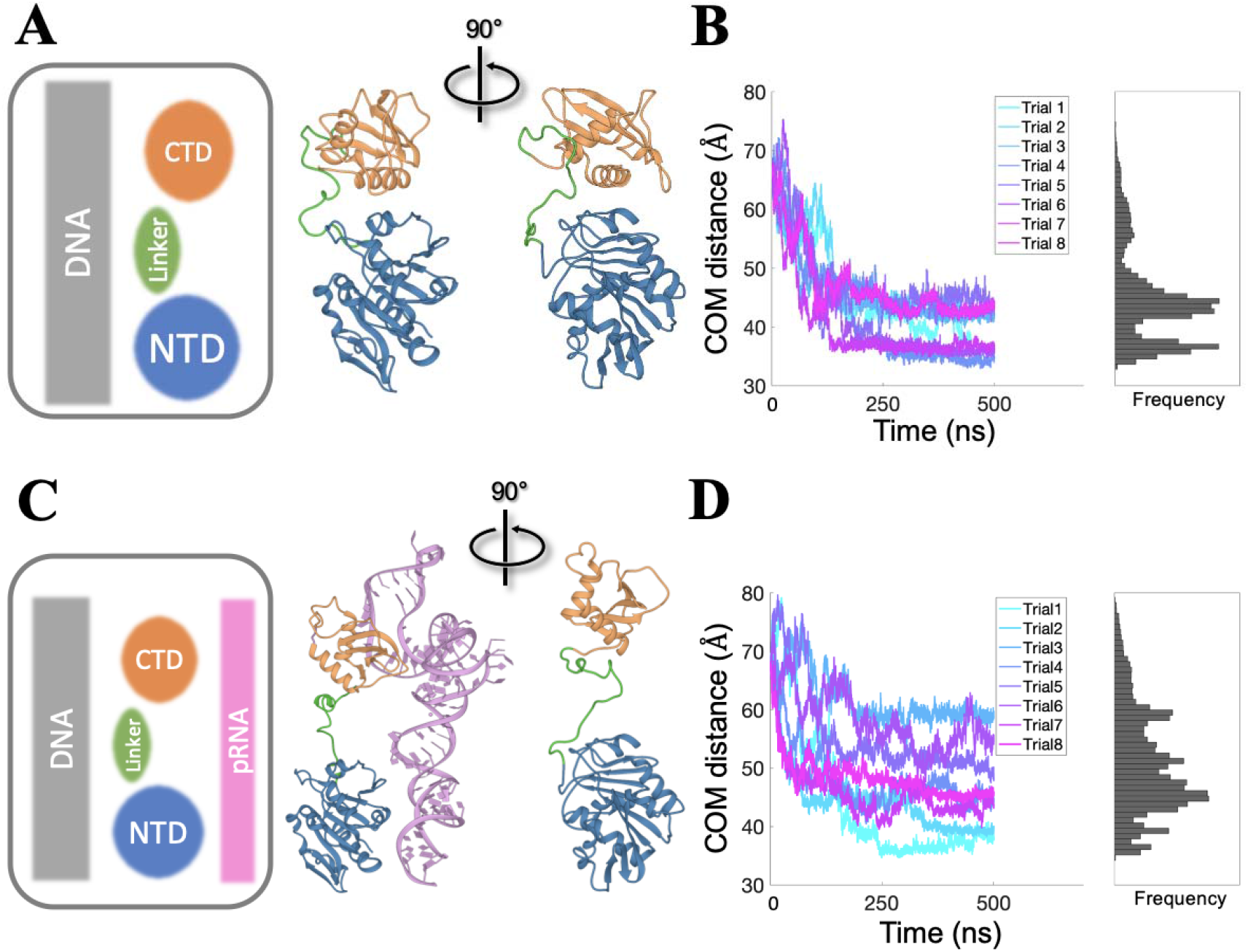
Investigating the dynamic interaction between NTD and CTD in the presence and the absence of pRNA with molecular dynamics (MD) simulations. (A) Simulation setup in the absence of pRNA and a snapshot of FL-gp16 in the compact mode. (B) Center-of-mass (COM) distances between NTD and CTD as a function of time during the simulations of FL-gp16 alone. (C) Simulation setup in the presence of pRNA and a snapshot of FL-gp16 in the extended mode. (D) COM distances between NTD and CTD as a function of time during the simulations of the FL-gp16/pRNA complex. Panel on the right shows the histogram of the sampled distances during the simulations.

Our MD simulations showed that a dominant population of sampled conformations has NTD-CTD COM separation of ~45 □ in which no direct contact between the domains was observed, hereafter referred to as the “extended” mode (Figure 4B). In three out of the eight trial simulations, the CTD was observed to be in touch with the NTD, hereafter referred to as the “compact” mode, and then bounce back to the extended mode. In the compact mode structure, the NTD-CTD separation is ~35 □ (Figure 4B) and the contact surface area is 754 □ ^2^. The binding interface involves an amphiphilic helix α2 from CTD and a short hydrophobic helix (residues 164-169) from NTD (Figure S6). The charged side of helix α2 faces NTD, and its electrostatic interaction with some charged residues on the NTD surface (K124 and D125) may draw the CTD towards the NTD. Upon the formation of this initial domain-domain contact, helix α2 rearranges itself to bury its hydrophobic surface inside the hydrophobic region on the NTD surface composed of V162, V163, F167, L168, F169, F170 and L172. A similar way of placing an amphiphilic helix in a hydrophobic environment is also found at the NTD-CTD interface of T4 gp17 ^8^, whereas the charge-charge interaction between NTD and CTD in T4 gp17 has not been confirmed. The only relevant mutagenesis study in which a Trp→Ala mutation inhibited DNA packaging and ATPase activity could provide only limited evidence for this NTD-CTD interaction ^8^.

Our simulations suggest that although possible, the compact mode is energetically less favorable, consistent with its abundance in several crystallographic structures of full-length viral motor proteins. However, assembly of these structures into oligomer structures, such as Sf6 gp2, would result in considerable steric clashes between the protein and DNA or adjacent subunits (Figure 3), implying that these NTD-CTD compact models might not represent their functional states but have instead resulted from crystal packing artifacts. Thus, modeling the full-length gp16 monomer alone in the absence of the other motor components is insufficient to picture a functional motor complex.

We next incorporated pRNA into the FL-gp16 model and probed the dynamic interactions between gp16-CTD and gp16-NTD in the presence of pRNA. We first docked the atomic structures of gp16-NTD and pRNA into the cryo-EM density map to estimate their relative positions (Figure S7A). We note that the pRNA density would be occupied by corresponding protein components in other viral packaging motors (Figure S8), consistent with the fact that pRNA is a unique component of the □29 motor. We then combined the gp16-NTD/pRNA, an MD-equilibrated gp16 linker, and a randomly oriented gp16-CTD to construct a gp16/pRNA binary complex model. Similar to the gp16-only system described above, eight MD simulations were performed starting from eight gp16/pRNA models with CTD in four different orientations and CTD-NTD separation of ~60 Å (Figure 4C; Movie S2). In the presence of pRNA, the NTD-CTD COM separation spanned 40 □ to over 60 □ during the simulations due to the interactions of CTD with the pRNA backbones (Figure 4D). The distance probability distribution peaked around 45 □, coinciding with the “extended” mode observed in the absence of pRNA. The space between the domains was occupied by the unstructured linker domain. At far separations, gp16-CTD makes contact with the three-way-junction (3WJ) of the pRNA. Several basic residues lining Site I of CTD were found engaged in stable electrostatic interactions with the negatively charged pRNA backbones (Figure 5B). We further inspected whether CTD has an orientation preference with respect to pRNA. Our simulation results showed that CTD preferably interacted with the pRNA through Site I in which both the major and minor binding grooves contributed to the binding (Figure S9). In addition, the linker domain appeared to provide the flexibility to orient CTD for optimal interaction with the pRNA. The other nucleic acid binding Site II is located on the other side of CTD facing the channel interior, but it is yet to be determined whether this second binding surface is utilized for DNA translocation.

**Figure 5.**
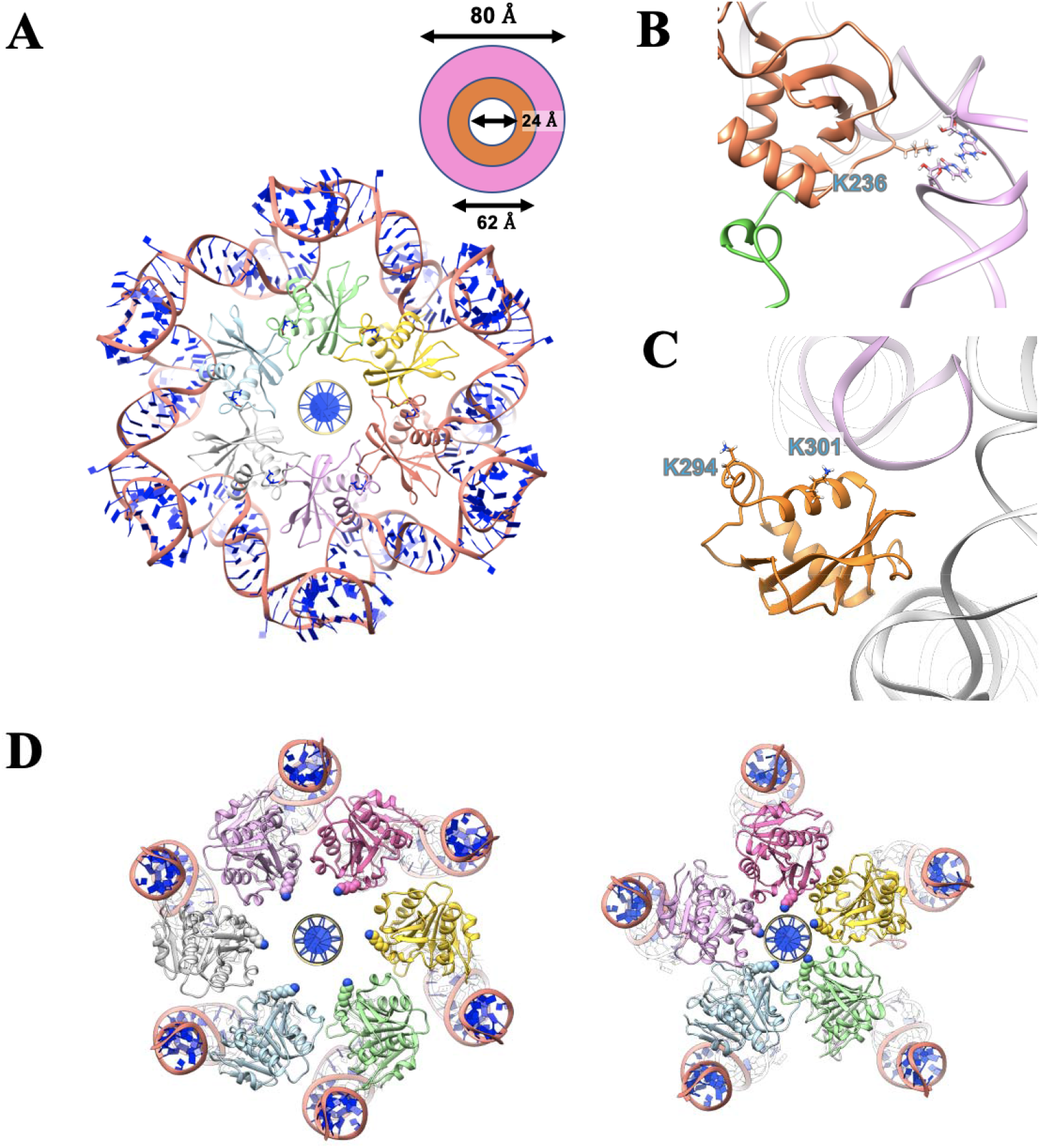
Construction of ring-shaped gp16 oligomers. (A) A hexameric ring of gp16-CTD and pRNA. Subunits are colored in green, yellow, red, purple, light blue and gray, respectively. pRNA bases are colored in blue and the backbone is colored in pink. A DNA molecule is placed at the center of the channel showing the relative size of the channel. (B) Interaction between the CTD (colored in orange) and the pRNA (colored in pink). The linker domain (partially shown) is colored in green. Interacting residues are labeled and shown as sticks. (C) Interaction between the CTD (colored in orange) and the adjacent pRNA (colored in gray). Interacting residues are labeled and shown as sticks. (D) A hexameric (left) and pentameric (right) ring of gp16-NTD. Similarly, subunits are colored in green, yellow, red, purple, light blue and gray (only in hexamer), respectively. K56 is shown as spheres in the color of the respective subunit.

### Interactions of gp16 with other motor components

Many previous studies have hinted potential interactions between the individual components of the □29 DNA packaging motor^16,18,38,39^, providing indirect evidence of how this motor complex assembles and operates. For example, the pRNA has been reported to bind to the N-terminus, i.e. the narrow end, of the connector ^25,39,40^. The 5’/3’ paired helical region of pRNA was revealed to be responsible for its interaction with gp16 ^38^. However, before the determination of an atomic structure of gp16-CTD, building a complete structural model of the motor complex has been impossible, although the oligomeric states of multiple individual components have been proposed and supported by a handful of biochemical and biophysical assays ^16,18,37,41–43^.

We constructed a complete structural model of the □29 genome packaging motor by integrating structural and experimental data from multiple sources (Figure 5, 6A, and S7B). We incorporated the connector structure into the gp16/pRNA binary complex model by docking the most recent atomic structure of the connector ^20^ into the cryo-EM density of the motor complex ^4^ to estimate their relative positions (Figure S7C). The ternary motor complex structure was then refined using stepwise MD simulations (details in Methodology) and the RMSD of protein backbone atoms from the initial structure was within 4 Å. The integrated structural model revealed interesting findings that would not be available from the structures of the individual motor components. pRNA was reported to form a hexameric ring structure through hand-in-hand interactions ^16^. In our model, gp16-CTD resides in-between two pRNA subunits interacting simultaneously with both RNA strands (Figure 5A). While gp16-CTD could assume various orientations, the basic residues such as K301 on helix α1 were found to always interact with an adjacent pRNA subunit (Figure 5C). Multiple interactions between gp16 and pRNA support the hypothesis that the pRNA ring facilitates the □29 motor assembly and operation by scaffolding gp16. While Site I orients towards pRNA subunits, the loop L2 between helix α1 and α2 extends to the interior of the motor channel. Basic residues such as K294 on L2 can bind the substrate DNA during translocation. Furthermore, the amphiphilic helix α2 is also exposed to gp16-NTD, providing a structural mechanism for potential inter-domain binding observed in simulations and crystallographic structures ^8,12^.

**Figure 6.**
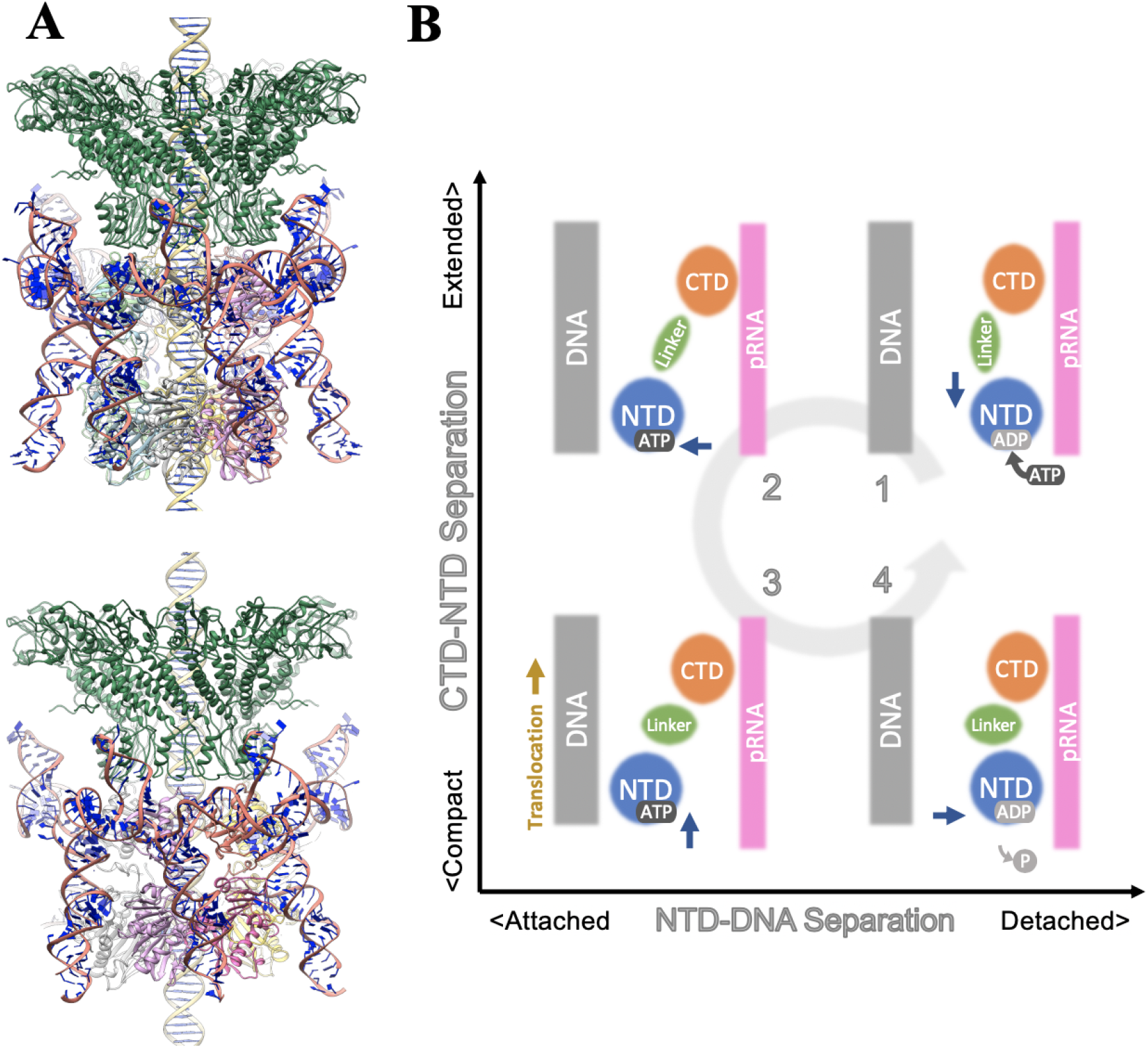
A hexameric structural model of the □29 DNA packaging motor complex and its implication for DNA translocation mechanism. (A) Snapshots of the motor complex with NTD in the “extended” mode (above) and the “compact” mode (below). The connector is colored in dark green. (B) Schematic of the inchworm revolving mechanism. The horizontal axis represents the NTD-DNA interaction: bound (attached; left) and unbound (detached; right) states. The vertical axis represents the NTD-CTD interaction: bound (compact; lower) and unbound (extended; upper) states. Motions of the NTD are indicated by blue arrows. Direction of the DNA translocation is indicated by yellow arrow.

Our hexameric gp16-NTD structure was constructed based on the hexameric HSV1 pUL15 ATPase ring. In addition to the previously identified DNA translocating loop containing R122 and K124 ^4^, K56 is also found situated in the interior of the motor channel (Figure 5D), suggesting that this positively charged residue may also be involved in mediating DNA interaction and translocation. The Walker residues G29, K30, S31, D118, E119, and the trans-acting arginine finger R146 are found at the interface of two adjacent gp16-NTDs, in agreement with a large body of mutagenesis and biochemical assays ^4,37,42,44^ We note that although there is a large discrepancy in the size of the two packaging motors: a diameter of ~150 □ for □29 versus ~225 □ for HSV1, the orientation of the ATPase domain relative to the motor channel appears to be conserved in the two packaging motors. However, we cannot exclude the possibility that the gp16 ATPase ring adopts a different structure or even changes its structure (e.g., diameter) during packaging. Importantly, given the scaffolding role of pRNA, an accurate description of the size of the gp16 ATPase ring will require high-resolution structural data of pRNA, especially the A-helix that extends to the NTD. While the structure and function of the 3WJ of pRNA have been studied extensively ^16–18,41^, the A-helix of pRNA was either missing or of very low resolution in previous cryo-EM studies ^18,22^.

Another important factor that may influence the gp16 ATPase ring size is the long-debated stoichiometry of pRNA. Both pentameric and hexameric pRNA oligomers have been proposed and supported by previous experimental data, and they would impose distinct constraints on the size of the ATPase ring. For the purpose of comparison, we have also constructed a □29 motor with pentameric stoichiometry by docking five pRNA and five FL-gp16 structural models into the cryo-EM density (Figure 5D, S10). The structure was then refined using molecular dynamics flexible fitting (MDFF) ^45^. As only low-resolution cryo-EM density (12 □) ^4^ is available, five-fold symmetrical restraints were imposed ^46^ throughout the refinement. Unsurprisingly, the size of the gp16-NTD ring decreases significantly from 40 □ in the hexameric model to 26 □ in the pentameric model (Figure 5D), which would suggest full contact of the substrate DNA with the motor channel during packaging. Nevertheless, our predicted critical intermolecular interactions between CTD and pRNA remain the same for both pentameric and hexameric stoichiometry: gp16-CTD is stabilized by pRNA that bridges gp16 and the connector, and anchoring gp16-CTD to the pRNA scaffold facilitates gp16’s transition between the “compact” and “extended” modes.

### Implication for DNA translocation mechanisms

Several translocation mechanisms have been proposed for ring-shaped biopolymer translocating biomotors. A hand-over-hand model was originally proposed to describe how the nucleic-acid-binding loops of a planar ATPase ring in helicases move along the translocation axis during the ATP hydrolysis cycle ^34,36^. Later, this model was extended to subunits in a non-planar ring moving along the helical axis to translocate ssDNA during DNA unwinding ^35,47,48^ or polypeptide chain during protein degradation ^49^. In line with the hand-over-hand model, a similar inchworm model in which a multi-domain ATPase undergoes conformational changes along the substrate translocation axis was proposed for the dsDNA packaging motor gp17 in bacteriophage T4 ^8,50^. Although a push-and-roll mechanism ^51,52^ has long been proposed for genome packaging in bacteriophages, previous biochemical and single-molecule experiments showed that this paradigm is not suitable for the □29 DNA packaging motor ^53^. To date, no direct structural evidence has been found supporting a rotary sequential model for the □29 packaging motor. Alternatively, a sequential revolving model, in which the substrate is pushed in a circular motion into the viral procapsid without rotation of any motor components, has also been proposed ^37,42,54–56^. In light of the recent structure of the herpesvirus terminase complex ^13^, a much larger size of the central channel (39 Å in diameter) than a B-form dsDNA (~20 Å in diameter) makes the sequential revolving model very attractive for the dsDNA packaging motors.

Our computational results showed that gp16 is highly dynamic with the CTD and NTD domains undergoing large displacements with respect to each other during the simulations, similar to T4 ^8^. Our complete structural model of the □29 motor complex further revealed direct interactions of gp16-CTD with the scaffold pRNA, which may help anchor gp16-CTD while allowing gp16-NTD to move during packaging. Together with the possible structural rearrangement of the gp16 ring, the transition between the “compact” and “extended” modes prompted us to propose a modified sequential revolving model, namely the inchworm revolving model, for □29 genome packaging (Figure 6B). In this model, gp16-CTD is anchored to the “stationary” pRNA, while the movement of gp16-NTD leads to transition of gp16 between the “compact” and the “extended” modes, which provides unidirectional forces to push the substrate DNA into the viral capsid. DNA translocation is likely mediated by a transient DNA binding surface in gp16-CTD and multiple DNA interacting sites in gp16-NTD, as identified in our modeled structure. Besides the movement along the DNA translocation direction, the conformational changes of gp16 also occur in the radial direction of the gp16 ring with the “attached” mode indicating the binding of DNA to gp16-NTD and the “detached” mode denoting the unbound state. As in the original revolving mechanism ^42,44^, the conformational changes of gp16 are coupled with ATP activities, and binding or hydrolysis of ATP shifts the binding affinity of gp16 for DNA and therefore sequentially activates individual gp16 subunits of the ATPase motor.

Schwartz *et al*. have shown that binding of ATP or γ-S-ATP (non-hydrolysable ATP) to gp16 significantly increases its binding affinity for DNA while ATP hydrolysis or adding an excess amount of ADP facilitates the release of DNA from gp16 ^44^. In our model (Figure 6B), ATP binding triggers gp16 transitioning to the “extended” mode along the translocation direction in which gp16-NTD moves away from gp16-CTD that stays stationary while anchored to pRNA (Step 1, upper right). Meanwhile, the gp16 ring is in its “attached” mode in which DNA is bound to gp16-NTD with high affinity (Step 2, upper left). Subsequently, ATP hydrolysis returns gp16 to the “compact” mode, push the gp16-bound DNA into the capsid (Step 3, lower left). Finally, DNA is released from gp16 as the bound ADP shifts gp16 to a low DNA binding affinity conformation (Step 4, lower right). During packaging, the anchoring of gp16-CTD to the pRNA scaffold is crucial so that the movement of gp16-NTD can be turned into a ratchet-like motion to drive DNA translocation.

Single-molecule experiments have provided valuable kinetic insights into how the viral packaging motors work ^57–59^. Temporary pauses and slips of DNA translocation observed in these experiments have been used to deduce the functional states and the coordination mechanisms of the ATPase rings ^60–62^. However, a one-way valve mechanism has also been proposed, in which besides the ATPase motor, the connector is also involved in controlling DNA translocation ^54,56,63^. Therefore, extra caution is required to interpret kinetic results in singe-molecule experiments as DNA packaging in □29 is likely a combinative process of pushing (ATPase) and valving (connector).

## Conclusion

We have performed an integrated study using X-ray structural determination and molecular modeling and simulations to elucidate the roles that the C-terminal domain of the □29 ATPase gp16 may play in motor assembly and genome packaging. Despite the lack of nuclease activity, the C-terminal domain is found to preserve a nucleic acid binding groove on the protein surface with an RNase H-like topology. Our structural comparison and MD simulations demonstrated that this surface site is better suited to the binding of pRNA than DNA. The C-terminal domain also possesses a transient DNA binding surface and may thus play a role in DNA translocation. Furthermore, our simulations highlighted a dynamic interplay between the N-terminal and C-terminal domains of gp16, which is dependent on the presence of pRNA. Finally, our computationally built and refined structural model of the complete □29 packaging motor complex revealed molecular details of the interactions between the C-terminal domain of gp16 and pRNA, which are essential for motor assembly and function. Taken together, our results suggested a refined translocation model for the □29 genome packaging motor, namely the inchworm revolving mechanism, which could account for a large body of previous structural and biochemical data. Our modified mechanistic model adds a new dimension of conformational dynamics to the original revolving mechanism, and also provides a mechanical explanation for the unidirectional translocation of DNA in the motor.

## Materials and Methods

### Cloning, expression, and purification of the C-terminal domain of gp16

The DNA fragment encoding the gp16-CTD (residues 209–332) was amplified by PCR and inserted into the LIC site of the vector pMCSG7 ^64^. The expressed proteins are fused with an N-terminal His tag. The gp16-CTD was expressed in *E. coli* B834 (DE3) cells overnight at 16°C to an optical density at 600 nm (OD600) of 2.2. Selenomethionine gp16-CTD was prepared as described previously ^65^. Protein expression was induced by the addition of isopropyl β-D-1-thiogalactopyranoside (IPTG) to a final concentration of 1.0 mM at an OD_600_ of 0.59, and cells were kept to grow overnight. Cells were harvested at 5,000 rpm, 4°C, 20 min, resuspended in a buffer containing 50 mM Tris-HCl (pH 8.5), 2 M NaCl, 10 mM imidazole and sonicated. Insoluble materials were sedimented by centrifugation (17,000 rpm, 4°C, 30 min). Protein was purified by Ni^2+^ affinity chromatography followed by size-exclusion chromatography on a Superdex 75 column (GE Healthcare) equilibrated in 20 mM Hepes pH 7.5, 250 mM NaCl, and 2 mM DTT.

### Crystallization

Crystals were obtained by sitting drop vapor diffusion in which 0.8 μL of a selenomethionine-labeled aliquot of gp16-CTD was mixed with an equal volume of reservoir solution containing 0.1 M Bis-Tris pH 6.5, 1.8 M ammonium sulfate and 2% PEG monoethyl ether 550 as a precipitant. Crystals grew after approximately 15 days and were flash-frozen directly from the drop in liquid nitrogen for data collection at 100 K.

### Structure determination and refinement

The X-ray diffraction data were collected at the Shanghai Synchrotron Radiation Facility (SSRF) beamline BL-17U1. Datasets were processed automatically by the program autoPROC ^66^. The crystal structure of the SeMet-labeled gp16-CTD was solved by single-wavelength anomalous dispersion (SAD) method with the program AutoSol in PHENIX ^67^. The structure was built manually using the program COOT ^68^, and the refinement was performed with the program PHENIX using noncrystallographic symmetry restraints. The model was improved by alternating cycles of refinement. The final refinement cycles included TLS refinement. The gp16-CTD crystals (residues 228-332) belong to space group P4122 with a = 42.68, b = 42.48, and c = 135.86 Å, which were solved to 2.32 Å resolution using single-wavelength anomalous dispersion data from a seleno-methionine substituted protein. Refinement statistics are summarized in Table 1.

**Table 1.**
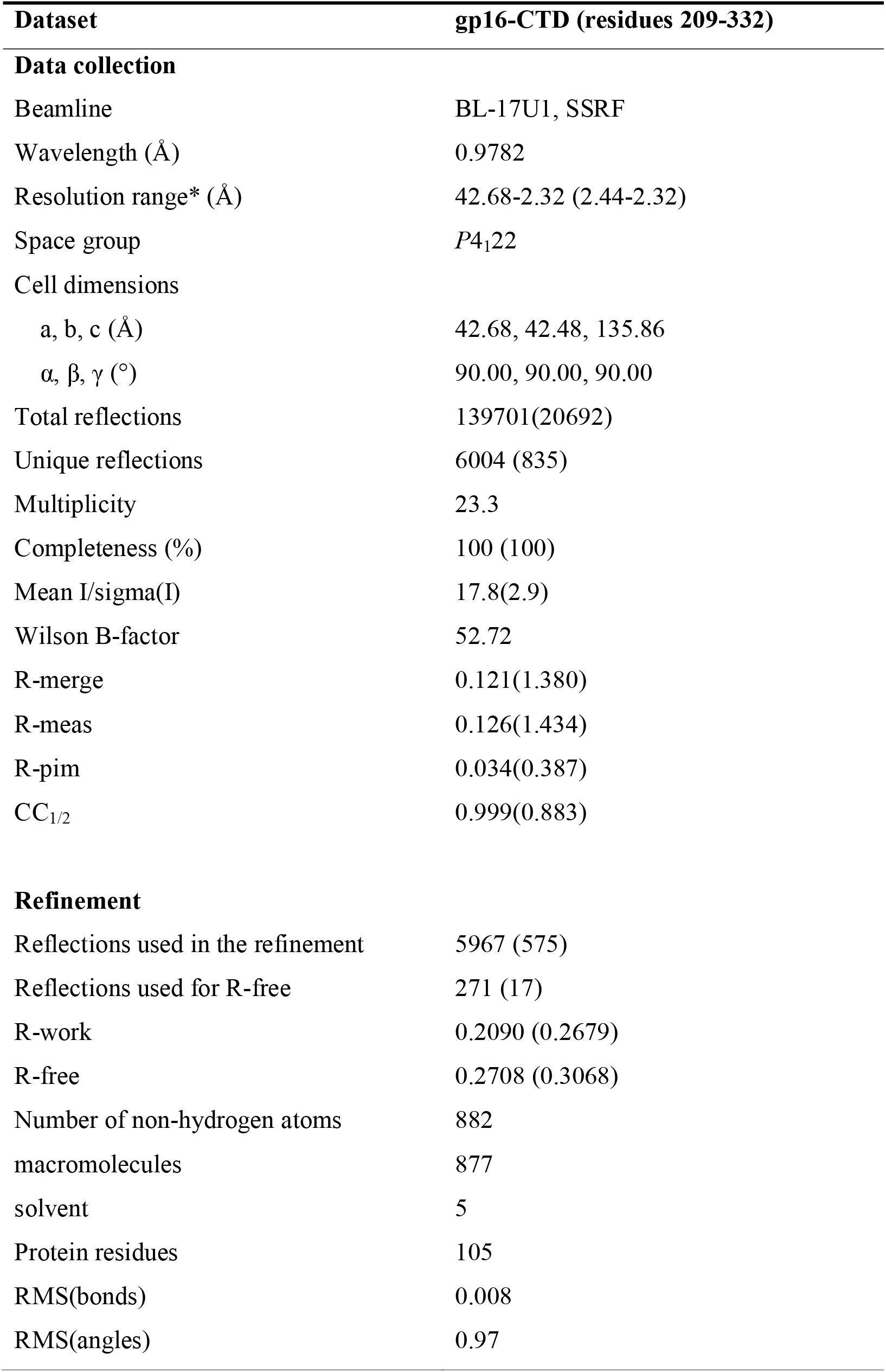
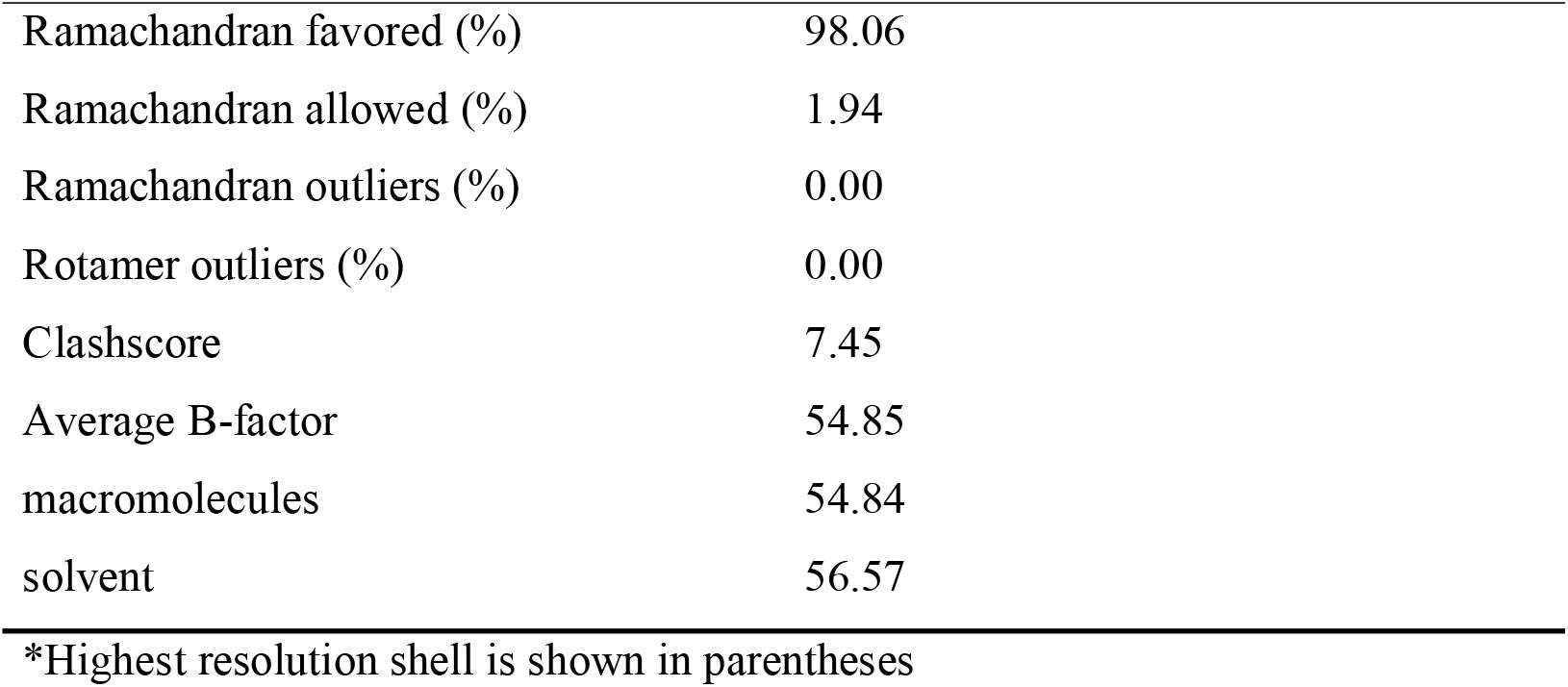
X-ray crystallography data collection and refinement statistics.

| **Dataset** | **gp16-CTD (residues 209-332)** |
| --- | --- |
| **Data collection** |  |
| Beamline | BL-17U1, SSRF |
| Wavelength (Å) | 0.9782 |
| Resolution range* (Å) | 42.68-2.32 (2.44-2.32) |
| Space group | *P*4<sub>1</sub>22 |
| Cell dimensions |  |
| a, b, c (Å) | 42.68, 42.48, 135.86 |
| $\alpha$, $\beta$, $\gamma$ (°) | 90.00, 90.00, 90.00 |
| Total reflections | 139701(20692) |
| Unique reflections | 6004 (835) |
| Multiplicity | 23.3 |
| Completeness (%) | 100 (100) |
| Mean I/sigma(I) | 17.8(2.9) |
| Wilson B-factor | 52.72 |
| R-merge | 0.121(1.380) |
| R-meas | 0.126(1.434) |
| R-pim | 0.034(0.387) |
| CC<sub>1/2</sub> | 0.999(0.883) |
| **Refinement** |  |
| Reflections used in the refinement | 5967 (575) |
| Reflections used for R-free | 271 (17) |
| R-work | 0.2090 (0.2679) |
| R-free | 0.2708 (0.3068) |
| Number of non-hydrogen atoms | 882 |
| macromolecules | 877 |
| solvent | 5 |
| Protein residues | 105 |
| RMS(bonds) | 0.008 |
| RMS(angles) | 0.97 |
| Ramachandran favored (%) | 98.06 |
| Ramachandran allowed (%) | 1.94 |
| Ramachandran outliers (%) | 0.00 |
| Rotamer outliers (%) | 0.00 |
| Clashscore | 7.45 |
| Average B-factor | 54.85 |
| macromolecules | 54.84 |
| solvent | 56.57 |

\*Highest resolution shell is shown in parentheses

### Molecular dynamics simulations

Molecular dynamics (MD) simulations were used to generate conformational ensembles of various structural components of the □29 DNA packaging motor, including gp16-CTD, linker domain of gp16, FL-gp16 with and without pRNA. We prepared the systems by solvating the solute in explicit TIP3P water molecules, with sodium and chloride ions at a final concentration of 0.15 M using the Gromacs *solvate* and *genion* tools ^69^. 1000 steps of steepest descent minimization were performed on the initial structure with all coordinates restrained except water. The system was further refined with a combination of steepest descent and conjugate gradient minimization approaches for another 5000 steps with protein backbone atoms harmonically restrained. After minimization, the system was heated to 310 K in a stepwise manner (50 K per 10 ps) using Berendsen thermostat with a coupling time constant of 0.5 ps followed by 500 ps equilibration stepwise releasing the harmonic restraints. Production runs were then carried out in NPT ensemble at 310 K and 1 atm, using velocity-rescaling thermostat ^70^ with a coupling time constant of 1 ps and Parrinello-Rahman barostat ^71^ with a coupling time constant of 2 ps. A timestep of 1 fs was used during annealing and 2 fs for equilibrium and production runs. Electrostatic interactions were calculated using the particle mesh Ewald sum method ^72^. A cutoff of 16 A□ was chosen for short-ranged van der Waals interactions. All the MD simulations were performed with Gromacs ^69^.

### Structural modeling and simulations of the full-length gp16

We built an atomic structure of the full-length pRNA based on a partial pRNA crystal structure (PDB: 4KZ2). We first completed the core part of the pRNA (base 25-95) by adding back missing nucleotides (nt) using Rosetta Stepwise application ^73^, which utilizes a Monte Carlo-based algorithm. The first and the last 25 nt (the extended part) were then modeled using Rosetta RNA 3D Structure Modeling application. Both the core and the extended parts were individually minimized and equilibrated using MD simulations with backbone phosphorus atom harmonically restrained before joining them together. Magnesium ions from the crystal structure were retained.

The linker domain (residues 198-227) of gp16 was built using Modeller ^74^ with the COMs of NTD and CTD separated by 60 Å. PsiPred ^32^ was used to predict the secondary structure of the linker domain as a reference. The linker domain was equilibrated in the solvent to generate an ensemble of structures with a length comparable to that inferred from the cryo-EM density (EMD-6560) (Figure S5). Ten linker domain structures were randomly chosen from the ensemble and placed in-between NTD and CTD. NTD was aligned to the HSV1 pUL15 oligomer structure (PDB: 6M5R), therefore having a relatively defined orientation. Four random orientations were taken for CTD (two with Site I facing the pRNA and two with Site II facing the pRNA). Residues 198-200 and 225-227 of the linker were then removed and re-modeled using Modeller in the presence of NTD and CTD to produce 100 structures of FL-gp16. Two structures with the lowest Discrete Optimized Protein Energy (DOPE) ^75^ were chosen for each orientation of the CTD as the initial structures (eight in total) for subsequent MD simulations. To mimic the environment in the motor complex, we also performed MD simulations of FL-gp16 in the presence of pRNA. The FL-gp16/pRNA complex structures were obtained by docking the atomic structures of FL-gp16 and pRNA into the cryo-EM density map. During the simulations, pRNA was restrained to its initial position, and a soft restraint of 2.5 kcal mol^−1^ Å^−2^ was applied if the COMs of NTD and CTD laterally deviated from their initial positions by >15 Å. Lateral displacement refers to the movement on the plane orthogonal to the NTD-CTD distance vector.

### Assembly and refinement of a complete structural model of the □29 motor complex

A hexameric oligomer of FL-gp16 was built by duplicating an equilibrated structure from the simulations of the gp16-pRNA complex and aligning all NTDs to those in the pUL15 oligomer (PDB: 6M5R). The dodecameric connector structure was obtained from previous studies (PDB: 6QX7). Three orientations of the connector were generated by rotating about its central channel axis by 10° each. The connector was placed next to the hexameric gp16-pRNA oligomer according to the cryo-EM density (EMD-6560) (Figure S7). Finally, a DNA molecule was built using 3DNA ^76^ and placed inside the central channel. For comparison, a pentameric motor complex was also built in a similar fashion. The only difference is that the pentameric FL-gp16 oligomer was built by duplicating the subunit based on five-fold symmetry upon NTD alignment. Both motor complexes were solvated as described in the MD simulation section and followed by simulation-based structural refinement.

The hexameric and pentameric systems consist of ~800,000 atoms and ~700,000 atoms, respectively. In the hexameric system, all Cα and phosphorus atoms were weakly restrained (1.0 kcal mol^−1^ Å^−2^) to their initial positions throughout the refinement simulation. After 100,000 steps of energy minimization, the system was brought to 300 K using Berendsen thermostat in 500 ps, followed by 1 ns of restrained MD simulation. The pentameric system was refined in alignment with the previous cryo-EM density (EMD-6560) using Molecular Dynamics Flexible Fitting (MDFF) ^45^. A five-fold symmetry has been enforced throughout the refinement ^46^. The force scaling was chosen to be 0.1. The cross-correlation coefficient (CCC) increased from 0.55 to 0.79.

## ACKNOWLEDGMENTS

This work was supported by National Natural Science Foundation of China (Grants no. 31770948, and 31570875) to S.O., the Special Open Fund of Key Laboratory of Experimental Marine Biology, Chinese Academy of Sciences (SKF2020NO1) to S.O., Marine Economic Development Special Fund of Fujian Province (FJHJF-L-2020-2) to S.O., and the High-level personnel introduction grant of Fujian Normal University (Z0210509) to S.O. The diffraction data were collected by S. O. at the beamline BL-17U1 of the Shanghai Synchrotron Radiation Facility (SSRF). X.C. acknowledges the start-up fund from the College of Pharmacy and the Discovery Themes at The Ohio State University. P.G.’s Sylvan G. Frank Endowed Chair position is funded by the Chen Foundation granted to the Ohio State University. No funding from P. G. has been used for this work.

## Conflict of Interest

P.G. is the consultant of Oxford Nanopore Technologies and Nanobio Delivery Pharmaceutical Co. Ltd, as well as the co-founder of Shenzhen P&Z Bio-medical Co. Ltd and its subsidiary US P&Z Biological Technology LLC, as well as ExonanoRNA, LLC and its subsidiary ExonanoRNA (Foshan) Biomedicine Co., Ltd.

