## Supplemental Info for "An integrative structural model of the full-length gp16 ATPase in bacteriophage phi29 DNA packaging motor"

**Movie Captions**

**Movie S1.** Molecular dynamics simulations of interaction between NTD and CTD in the absence of pRNA. Coloring of the protein domains was described in Figure 4. The full-length gp16 was shown to assemble a compact mode at the end of the trajectory.

**Movie S2.** Molecular dynamics simulations of interaction between NTD and CTD in the presence of pRNA. Coloring of the protein domains was described in Figure 4. The full-length gp16 was shown to assemble an extended mode at the end of the trajectory.

**Table S1.** Structural homology between ϕ29 packaging ATPase (PDB: 5HD9) and other viral packaging ATPase.

| Protein | PDB | Pruned Cα pairs | RMSD (Å) |
|---|---|---|---|
| ϕ29 gp16 (homology model) | | 139 | 4.7 |
| HSV1 pUL15 | 6M5R | 117 | 5.2 |
| T4 gp17 | 3CPE | 153 | 4.5 |
| Sf6 gp2 | 4IEE | 161 | 4.4 |
| FtsK | 2IUU | 54 | 5.7 |
| ϕ12 P4 | 1W44 | 51 | 5.7 |

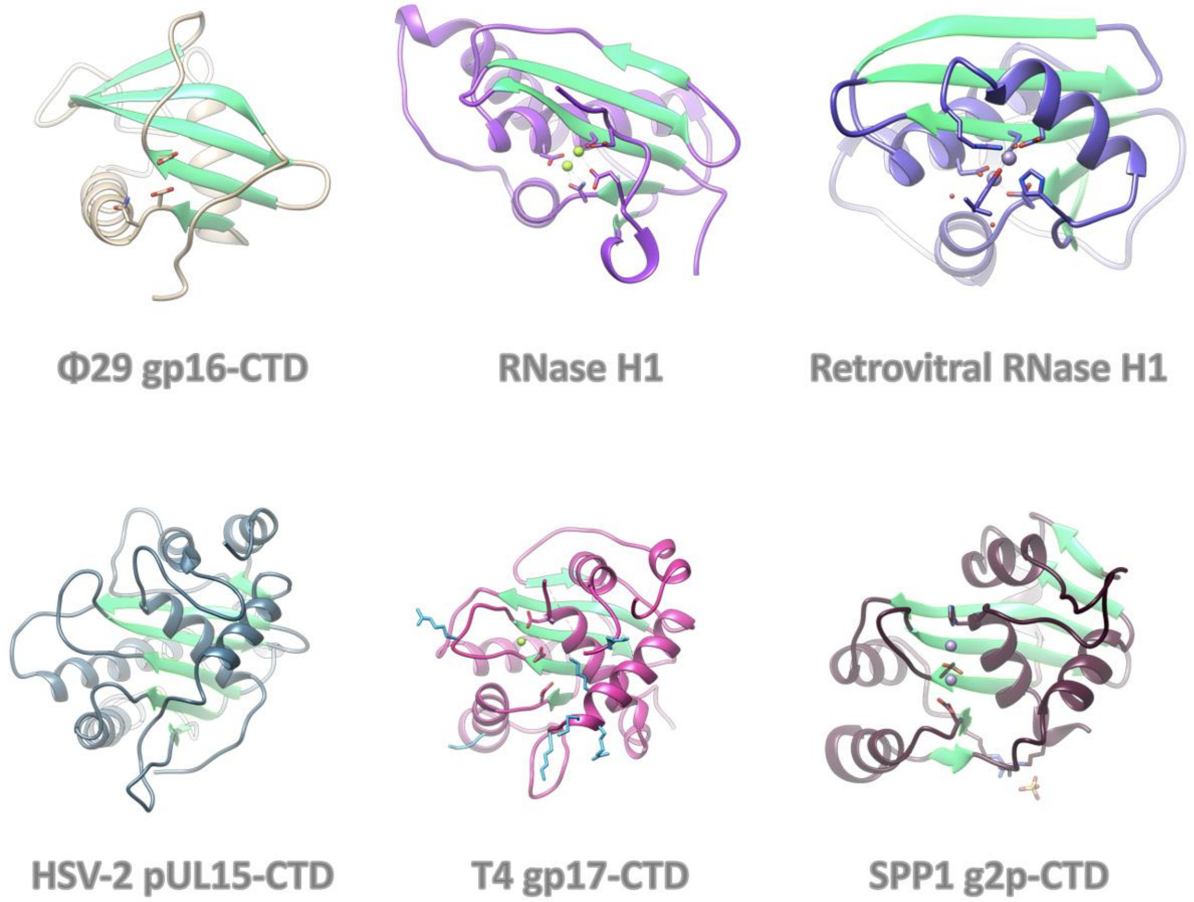


**Figure S1.** Comparison of gp16-CTD with RNase H domains and CTDs of other viral packaging motors. Catalytic residues are shown as sticks. The crystallographic magnesium ions (if any) are shown as spheres. For T4 gp17, residues proposed to interact with the substrate DNA are colored in cyan, which constitute a nucleic acid binding surface.

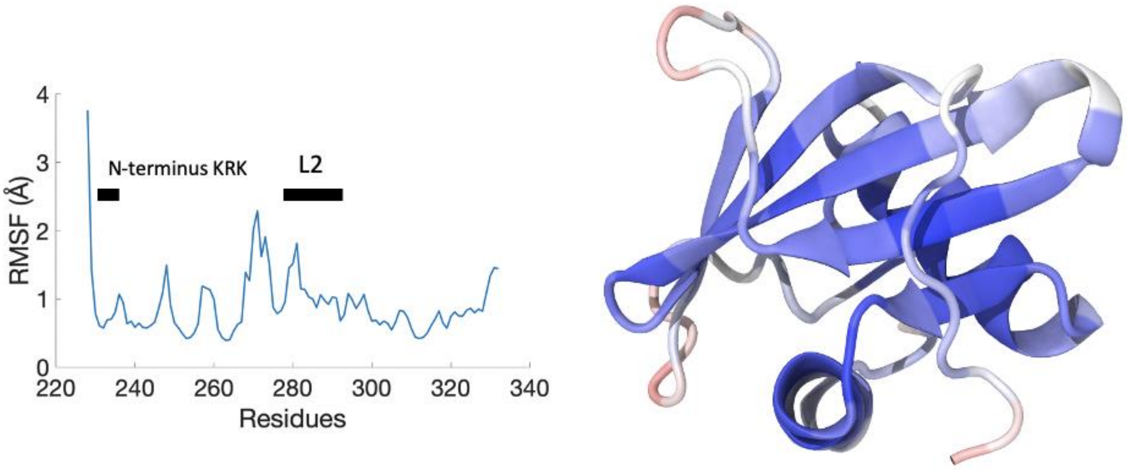


**Figure S2.** The root-mean-square-fluctuation (RMSF) profile of gp16-CTD in the MD simulation (left); and B-factor values mapped onto the gp16-CTD structure (right).

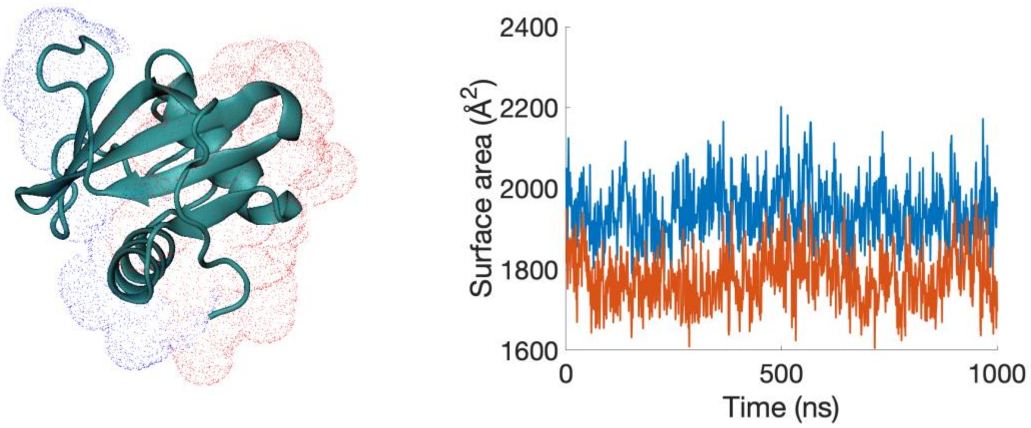


**Figure S3.** Solvent-accessible surface area (SASA) of gp16-CTD (left). SASAs for the major and minor binding surfaces as a function of time during the MD simulation of gp16-CTD (right). Major and minor binding surfaces are colored in blue and red, respectively.

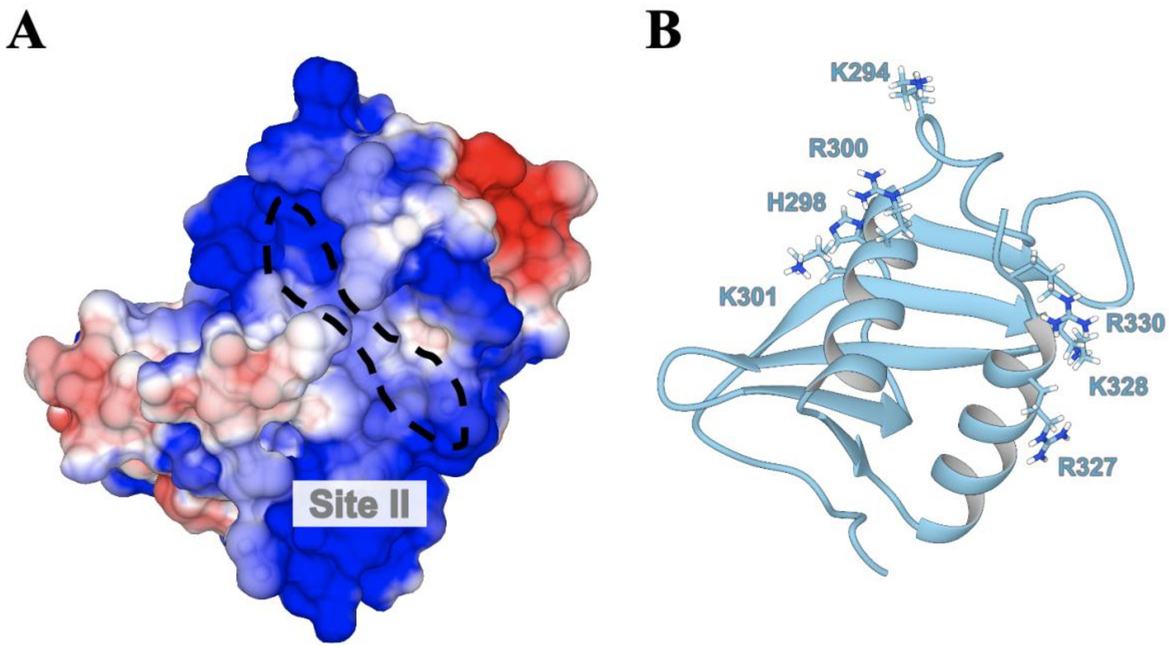


**Figure S4.** (A) Electrostatic surface of gp16-CTD with site II indicated by dashed lines. (B) Positively charged residues bordering site II on the surface are labeled and shown as sticks.

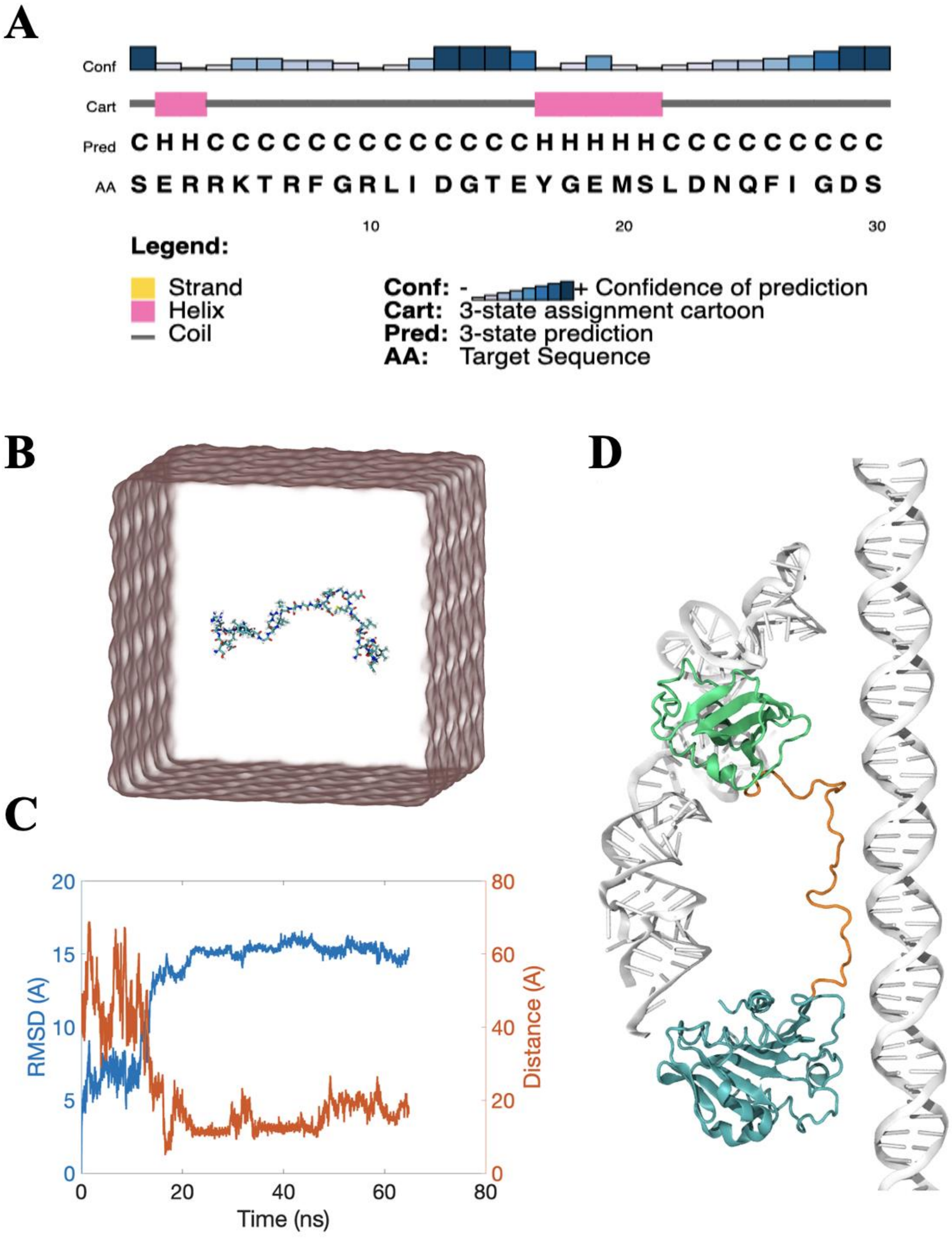


**Figure S5.** (A) Secondary structure prediction of the linker domain of gp16. (B) Simulation setup of the linker domain as an extended peptide chain in aqueous solution. (C) RMSD (blue) and end-to-end distances (red) of the linker domain as a function of time during the simulation. (D) Linker domain (orange) added in-between NTD (cyan) and CTD (green) of gp16 with the pRNA and DNA (silver) shown as references.

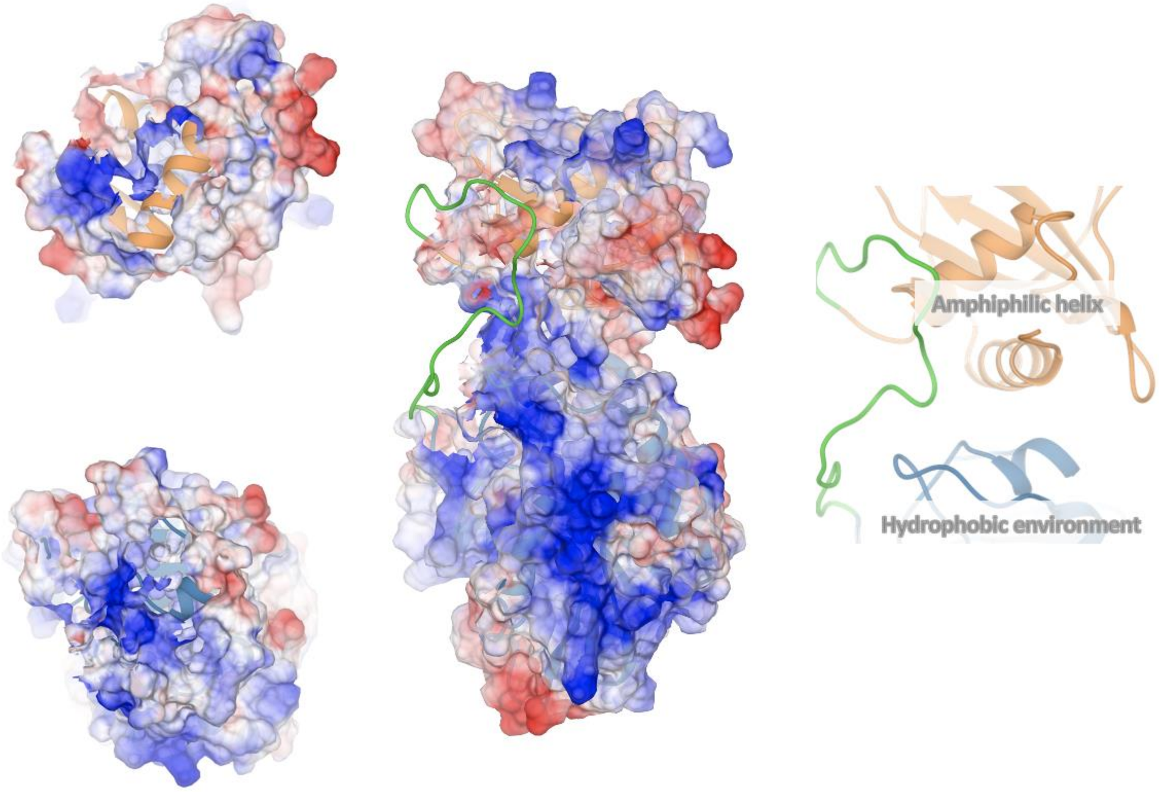


**Figure S6.** A snapshot from the MD simulation showing the NTD-CTD interaction interface. The linker domain is colored in green. Electrostatic properties are shown on the protein surface. Positive and negative charges are colored in blue and red respectively. The right panel shows a cartoon view of the interaction interface. CTD, linker domain and NTD are colored in orange, green and blue, respectively.

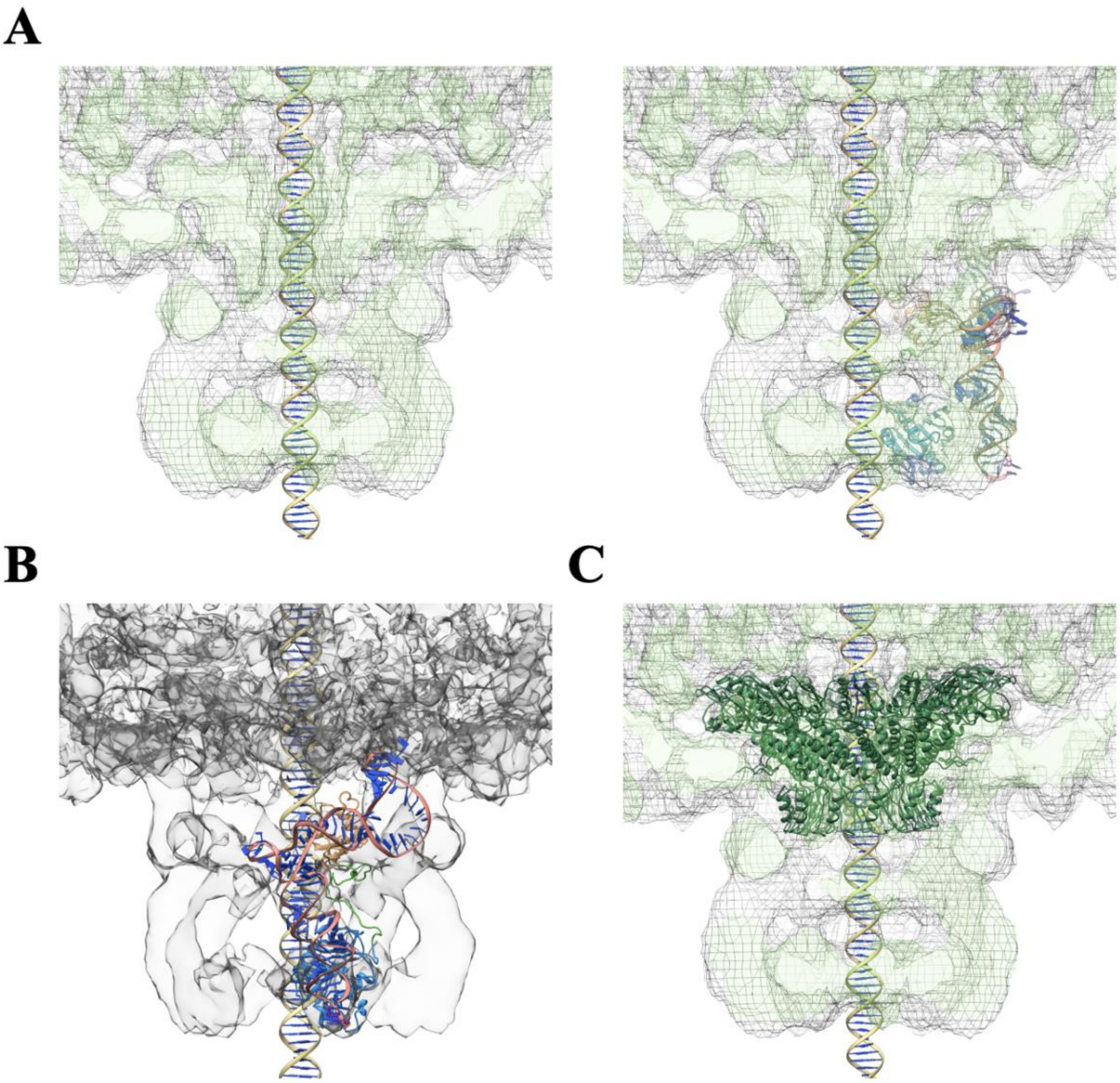


**Figure S7.** (A) Cross-section of the cryo-EM density map (EMD-6560) (left) and docking of a gp16-pRNA complex structure into the density (right). (B) A front view of a gp16-pRNA complex structure docked into the density. (C) Docking of the connector protein into the density.

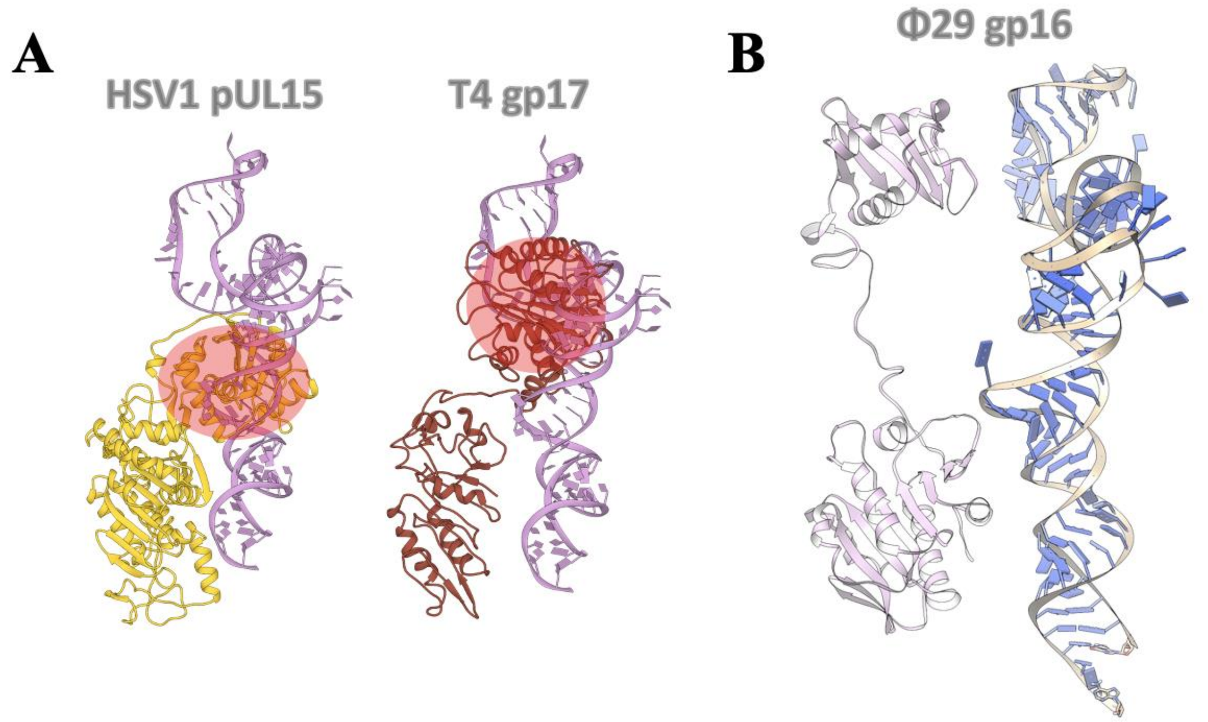


**Figure S8.** (A) Introducing pRNA to other viral packaging motor proteins induces steric clashes. (B) Placement of pRNA beside a full-length gp16 structural model according to the cryo-EM density (EMD-6560).

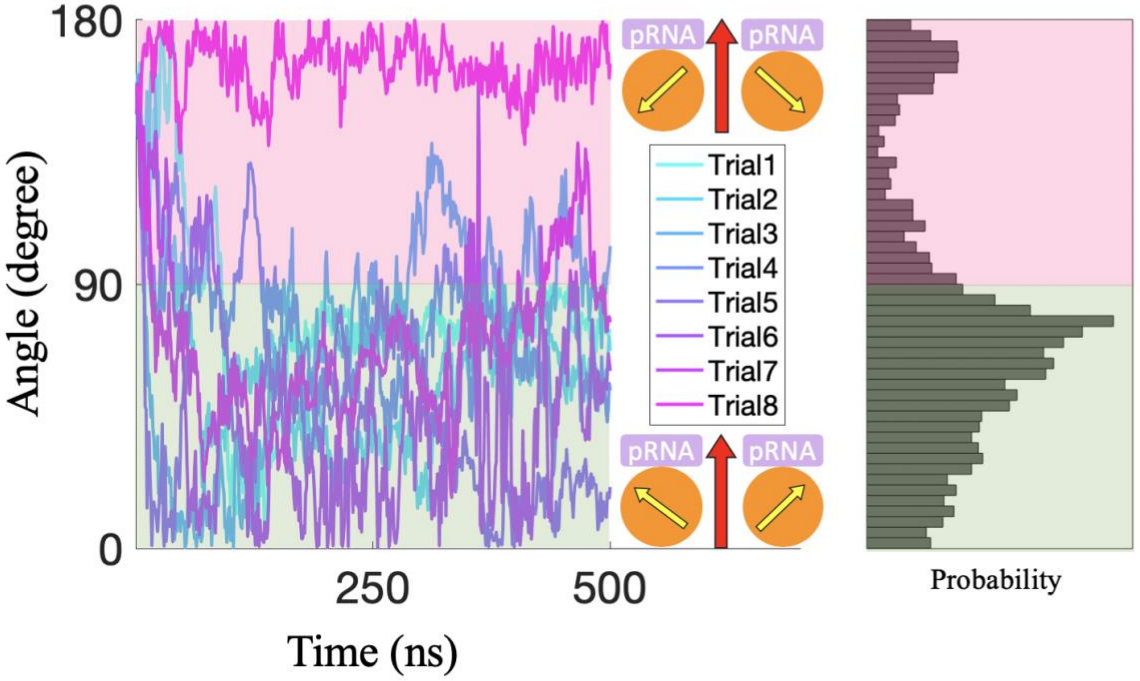


**Figure S9.** The orientation of gp16-CTD with respect to pRNA as a function of time during the MD simulations. The yellow arrow points from Site II to Site I of the gp16-CTD. The angle measures the deviation of the yellow arrow from the reference vector (red arrow) that points from the COM of gp16-CTD to that of pRNA . Therefore, an angle from 0° to 90° (shaded in green) indicates that Site I faces pRNA whereas an angle from 91° to 180° (shaded in red) indicates that Site II faces pRNA. The right panel is the histogram of the sampled orientations.

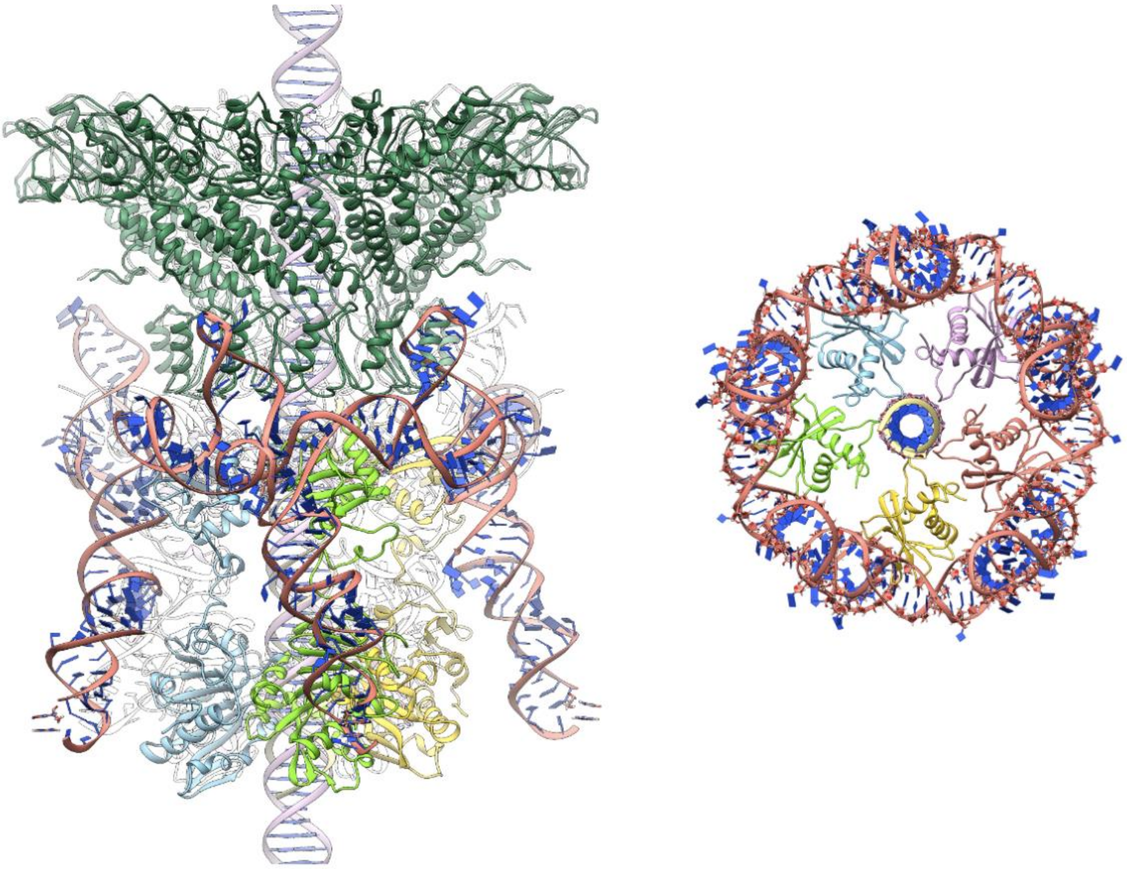

**Figure S10.** A hexameric structural model of the ϕ29 DNA packaging motor complex in a peripheral view (left) and a CTD-pRNA hexameric ring structure viewed from the capsid inside (right). Subunits are colored in green, yellow, red, purple, and light blue, respectively. The connector is colored in dark green. pRNA bases are colored in blue and the backbone is colored in pink. A DNA molecule is placed at the center of the channel showing the relative size of the channel.
